# Wearable sweat nicotine biosensor based on a nicotine oxidoreductase and its natural electron acceptor

**DOI:** 10.1101/2025.10.31.685570

**Authors:** Zhehao Zhang, Abdurrahman Addokhi, Kristin A. Hughes, Hao Zang, Maria Del Carmen Piqueras, Uroš Kuzmanović, Roger Charles, Leslie Leung, Catherine M. Klapperich, Karen N. Allen, Mark W. Grinstaff, James E. Galagan

## Abstract

The advent of wearable biosensors is empowering clinical and lifestyle decision making. Nicotine is a drug of significant negative impact on public health given the prevalence of smoking and vaping. Yet noninvasive, continuous monitoring of nicotine is challenging due to lack of specific and sensitive biosensing elements. Here, we report a wearable highly sensitive nicotine biosensor and demonstrate on-body deployment of the sensor in a first-in-human study. The biosensor comprises nicotine oxidoreductase (NicA2) with its natural cytochrome c electron acceptor (CycN) as mediator: both proteins play a key role in the nicotine-degrading bacterium, *Pseudomonas putida S16*. The biosensor detects nicotine over four-orders of magnitude (0.1-100 μM) with nanomolar sensitivity (LOD ∼33.6 nM), performs in the physiological pH range (pH 6-9), and is highly selective against common interferants and the major human nicotine metabolite, cotinine. Incorporation of the biosensor in a custom wearable device enables real-time measurement of nicotine from locally induced sweat on volunteer subjects and demonstrates higher analytical accuracy than gold-standard mass spectrometry.

## Introduction

Wearable devices are leading the next paradigm shift of healthcare, and real-time monitoring of physiology is key to this transition in patient care [1–9]. However, with the exception of continuous glucose monitors (CGM) nearly all existing wearable devices collect only physical parameters of the body such as heart rate and activity data through electrical, mechanical or optical sensors [10]. Current solutions for sensing chemical and molecular biomarkers, by contrast, typically involve invasive sampling of blood and lab instruments that are expensive, difficult to operate, not portable, and not translatable to point-of-care or wearables [11]. To fully reap the benefits of wearable health devices, biochemical sensors must be brought in contact with an appropriate biofluid, such as sweat, in a non-invasive fashion.

Nicotine is a naturally occurring plant alkaloid produced by tobacco species e.g., *Nicotiana tabacum*. It is the main addictive component of tobacco products and e-cigarettes and is a drug that negatively impacts public health [12–14]. In the body, nicotine binds to nicotinic acetylcholine receptors (nAChRs) of brain neurons and triggers a series of neurological events that lead to nicotine dependence [15]. Tobacco is the leading cause of preventable deaths worldwide and the addictiveness of nicotine is the primary reason for tobacco abuse [15, 16]. In recent years, electronic nicotine delivery devices such as e-cigarettes have emerged as an alternative to tobacco, but these can be equally harmful to lung and brain development [16].

Nicotine monitoring can be used to minimize nicotine exposure and help people quit smoking. Current nicotine monitoring and protection measures rely primarily on population-level surveys to keep track of smoking patterns and enforcing bans on smoking indoors and at public places. Unfortunately, such measures require considerable input and compliance from nicotine users and suffer a lack of evidence-based approaches. The gold standard for nicotine analysis is mass spectrometry (GC/LC-MS) [17, 18] which, although highly specific and sensitive, requires complex instrumentation, trained personnel, and long turnaround times, and thus is not suitable for public health applications. Wearable nicotine biosensors are a promising solution for detecting nicotine exposure and monitoring nicotine both at individual and population scale due to their compact size, fast analysis and cost-effectiveness. Another potential application is smoking cessation. Smoking cessation programs typically employ nicotine-replacement therapies (NRTs), however, efficacy is often hindered by the genetic and behavioral heterogeneity of smokers [19]. Monitoring nicotine as a biomarker could provide a personalized approach to account for such heterogeneity and improve the outcome of treatment. While a number of nicotine biosensors have been developed over the years [20–31], existing sensors generally lack specificity and sensitivity and have not yet been validated in real-time nicotine analysis in body fluids.

Electrochemical biosensors using enzymes as biorecognition elements represent the most advanced and commercially successful biosensors owing to the immense global market for glucose monitoring in diabetes management and the success of the enzyme-based continuous glucose monitor [32]. Compared to biosensors based on affinity, such as those dependent on antibodies, aptamers, or molecularly imprinted polymers (MIPs) which are difficult to regenerate after analyte binding, enzyme-based biosensors enable continuous monitoring owing to the catalytic nature of enzymes, their cost effectiveness, and the simplicity of the device. Enzymatic electrochemical sensors function by measuring the transfer of electrons generated by an enzyme-catalyzed redox reaction [33–37]. A central challenge is the electron transfer between the enzyme and the electrode due to the shielding of the redox centers of enzymes. To overcome the challenge, three generations of sensors have been developed based on different electron-transfer mechanisms, namely, the O_2_/H_2_O_2_ system, mediator-based systems, and direct electron transfer systems. Among these, the use of mediators presents a promising strategy for improving electron transfer, when the mediator can efficiently shuttle electrons between the enzyme and the electrode [38]. Although many mediator-based biosensors have been developed, they typically rely on artificial redox species such as ferrocenes, methylene blue, and transition-metal complexes [35, 39]. On the other hand, biological systems employ natural electron acceptors such as cytochromes, ferredoxins, azurins, etc. often co-evolved with the redox enzyme to enable specific and efficient electron flow of the metabolic pathway [40]. Despite the ubiquity and promise of these natural mediators, they have rarely, if ever, been applied in biosensors with their native enzyme partners.

Herein, we report the development of a highly specific nicotine biosensor with nanomolar sensitivity, derived from microbial biorecognition elements, and the non-invasive deployment of the sensor in a wearable device for real-time, continuous monitoring of nicotine from sweat. The biosensor is based on a nicotine oxidoreductase (NicA2) with its natural cytochrome c electron acceptor (CycN) as mediator. In a separate manuscript [41] we report the identification of NicA2 through a genomic screen we developed and applied to the nicotine-degrading bacteria *Pseudomonas putida S16* [42], and the subsequent development of first-generation prototype electrochemical sensors based solely on NicA2. In this manuscript we describe the development of a second-generation nicotine sensor based on both NicA2 and CycN. The sensor demonstrates the first use of the natural cognate electron acceptor with a redox enzyme to dramatically improve biosensor performance. The NicA2/CycN sensor demonstrates 1-2 orders of magnitude improvement in sensitivity compared to the first-generation sensors, sufficient to detect the full range of nicotine levels in human sweat. We deploy the biosensor on volunteer subjects in a custom wearable device for real-time measurement of nicotine from sweat and demonstrate higher analytical accuracy than mass spectrometry in classifying nicotine users from non-users. The findings demonstrate the effectiveness of the nicotine sensor for real-time detection of nicotine in sweat and highlight the potential for wearable devices to monitor chemical biomarkers with accuracy equivalent to gold-standard method.

## Results

In a separate manuscript, we describe the development of prototype sensors developed from WT NicA2 and multiple variants based on first-generation sensor design where electrons are transferred from enzyme-bound flavin adenine dinucleotide (FAD) to molecular oxygen (O_2_) to produce hydrogen peroxide (H_2_O_2_) that is subsequently oxidized or reduced at an electrode surface [41]. However, first generation sensors are limited by the efficiency with which O_2_ interacts with enzyme-bound FAD, and the diffusion of H_2_O_2_ to the electrode surface [38]. To address this limitation, second generation sensors use mediators to more efficiently shuttle electrons from the enzyme to the electrode [38]. Whereas many mediator-based biosensors have been developed, they are typically based on artificial redox species [35, 39]. In cells, by contrast, the regeneration of enzyme-bound co-factors is accomplished by electron transfer with a cognate natural electron acceptor often co-evolved with the redox enzyme to enable specific and efficient electron flow [43]. Cytochromes are a common natural electron acceptor for redox enzymes, and the use of cytochromes as mediators for second generation biosensors has been explored [44, 45]. However, the use of a naturally co-evolved cytochrome as a mediator for its cognate redox enzyme has not been previously reported.

### Identification of CycN using genomic screening

We previously screened the bacterium *Pseudomonas putida S16* (*P. putida S16*) known to metabolize nicotine [46] and identified a gene cluster that was significantly up-regulated in the presence of nicotine that we termed the nicotine responsive genome island (NRGI) [41]. Through this approach, we identified the previously reported gene for nicotine oxidoreductase (NicA2) that was up-regulated by 16-fold in the presence of nicotine [41]. The same cluster also includes a gene encoding a c-type cytochrome (CycN) co-expressed with NicA2 (15-fold up-regulated in response to nicotine) located directly downstream of the gene for NicA2. CycN was recently characterized as a natural electron acceptor of NicA2 [47, 48] and has been shown to be able to re-oxidize NicA2 up to 10,000-fold faster than oxygen in biochemical assays [47].

To explore the relationship of NicA2 to other oxidases and cytochrome-dependent dehydrogenases in the flavin amine oxidase superfamily, we performed a bioinformatic analysis of sequences homologous to NicA2. To identify sets of highly similar sequences, a sequence similarity network (SSN) was calculated using UniRef 50 (Figure 1A). Using an e-value cutoff of e^-5^ and an alignment score of 67 allowed clustering of nodes sharing >35% sequence identity with NicA2 (Figure 1B). To identify protein families that frequently co-occur in the same operon as homologs of NicA2, the sequence of NicA2 and 170 sequences in 25 connected nodes in the SSN were used as query sequences to calculate a genome neighborhood network (GNN; Figure 1C). Analysis of the GNN and corresponding genome neighborhood diagrams revealed dozens of homologs of NicA2 and CycN which co-locate within the genome neighborhood, in most cases with CycN encoded directly downstream of NicA2 as in *P. putida* (Figure 1D). Most strikingly, we identified multiple cases of gene fusions between homologs of NicA2 and CycN (Figure 1E). Together, these results suggest that NicA2 and CycN co-evolved to function together in nicotine degradation, leading to more efficient electron transfer. We therefore studied the potential of using CycN as a mediator for NicA2 in a second-generation sensor to increase electron flow and thus sensitivity.

**Figure 1.**
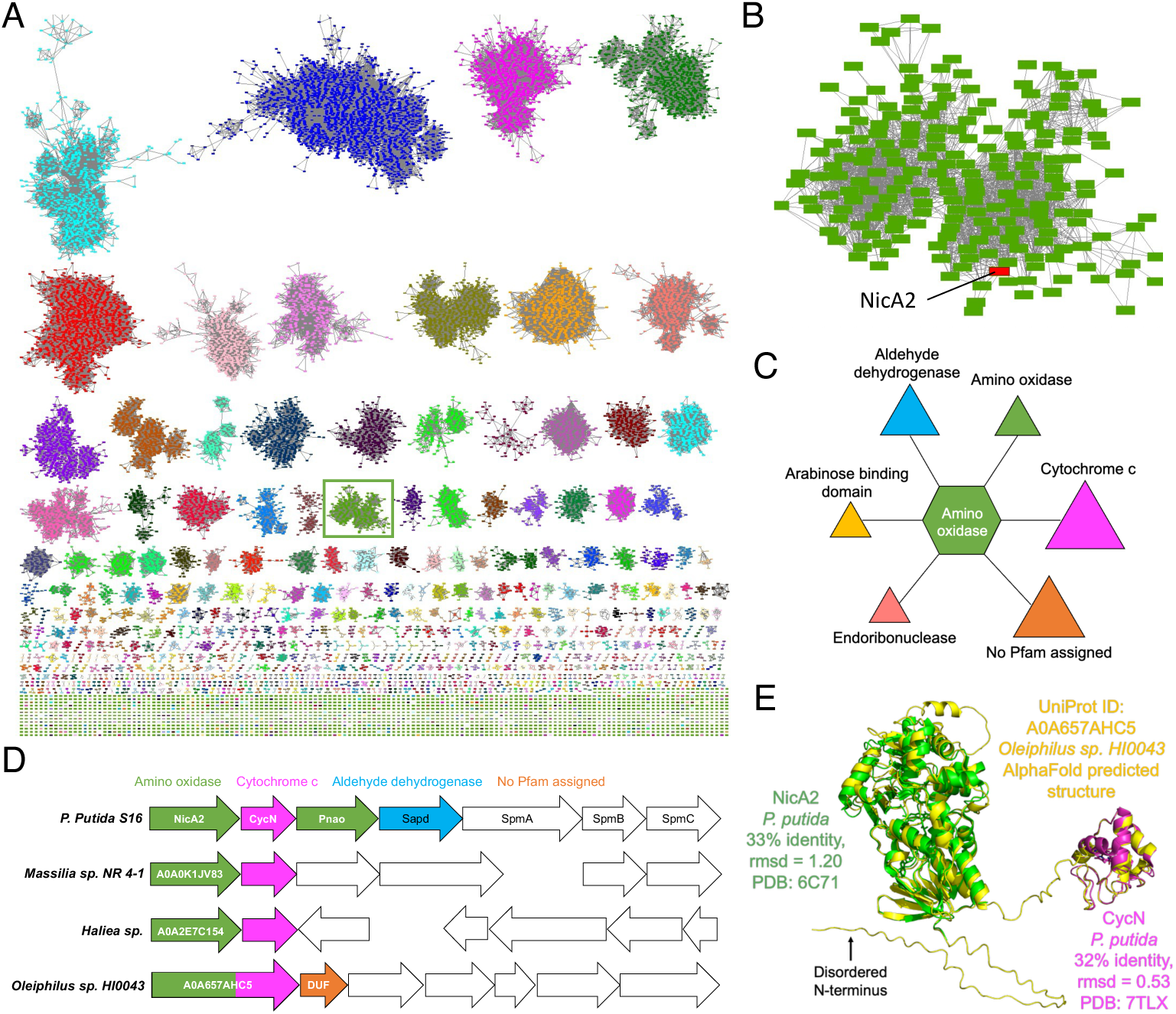
Genomic analysis suggests NicA2 and CycN have co-evolved to catalyze nicotine degradation. **(A)** Sequence similarity network (SSN) for the Flavin containing amine oxidoreductase family, Pfam PF01593 (short name, amino_oxidase). Cluster containing NicA2 (F8G0P2) is boxed in green. Parameters for SSN: UniRef50, E-Value = 5, Sequence Length = 200-1000, Alignment Score = 67 **(B)** Expanded view of the cluster containing NicA2 (highlighted) **(C)** Genome neighborhood network (GNN) for 170 sequences from the NicA2 cluster (Figure 1B) of the SSN. **(D)** Genome neighborhood diagrams for genes encoding NicA2 and NicA2 homologs (UniProt IDs provided) including a downstream cytochrome c (Pfam PF00034). **(E)** Predicted structure of the fusion protein A0A657AHC5 by AlphaFold 3 and overlaid with the experimental structures of NicA2 and CycN.

### Electrochemical characterization of CycN

To characterize the electrochemical properties of CycN, we expressed and purified the recombinant protein in *E. coli* and performed cyclic voltammetry (CV). We used a gold electrode modified with self-assembled monolayers (SAM) known to enable direct electron transfer for other cytochrome c proteins [49]. Cytochrome c proteins enable electron transfer through the reversible oxidation and reduction of the central ferrous/ferric iron (Fe^2+^/Fe^3+^) of the heme prosthetic group (Figure 2A) [50]:

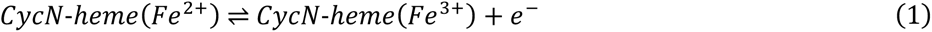

**Figure 2.**
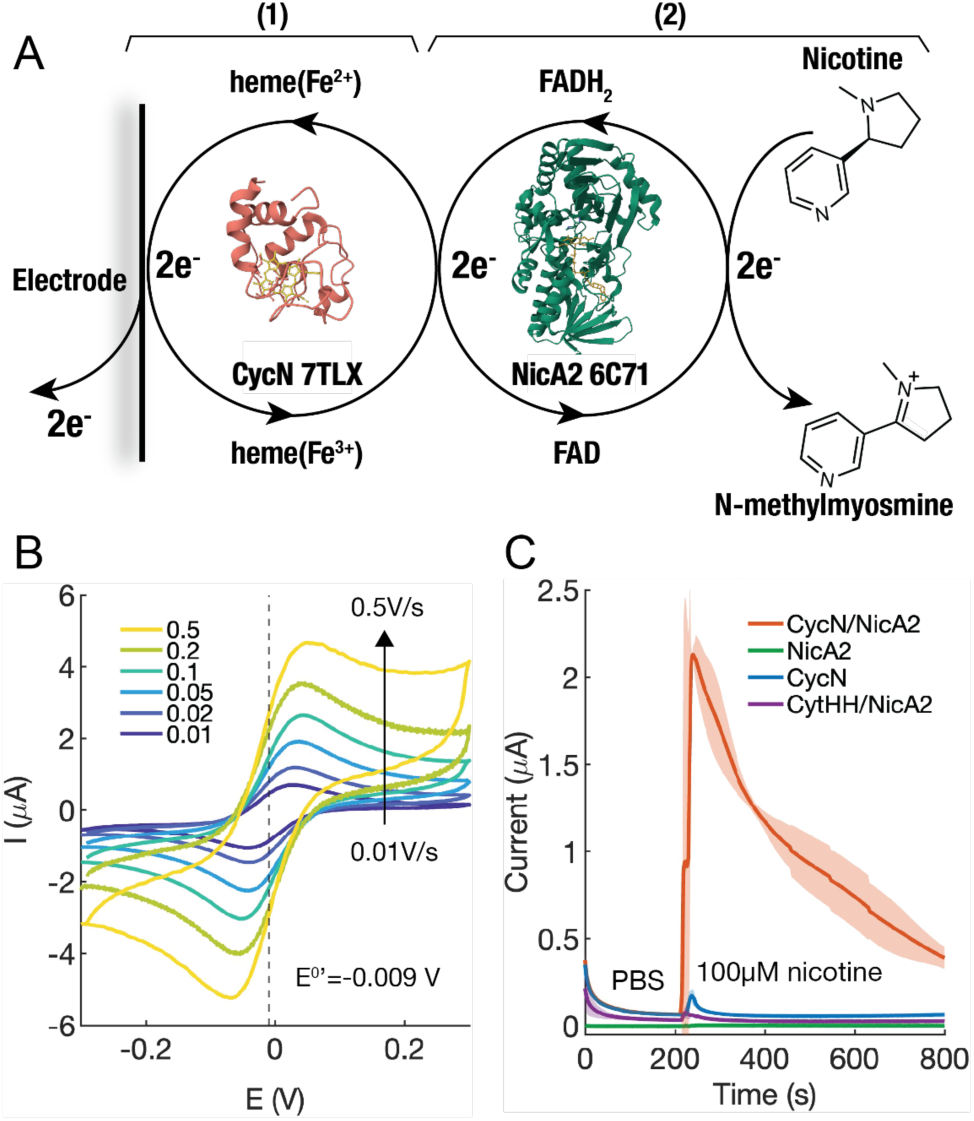
Electrochemical characterization of CycN confirms its redox activity and capability for mediating NicA2 catalysis. **(A)** Reaction steps involved in the CycN-coupled NicA2 catalysis. (1) Redox reactions of CycN. (2) Oxidation of NicA2 by CycN and enzyme reaction. **(B)** CycN shows stable and reversible redox behavior on gold electrode modified with self-assembled monolayers (SAM). The formal reduction potential (E^0**′**^) of CycN is −9 mV vs screen-printed silver reference electrode. PBS buffer is used as the electrolyte. [CycN] = 100 μM. Scan rate = 0.01 to 0.5 V/s. **(C)** CycN is capable of mediating NicA2 catalysis upon adding nicotine to the buffer, whereas neither NicA2 nor CycN alone respond to nicotine. No nicotine response was detected when NicA2 was co-incubated with another cytochrome c (*horse-heart cytochrome c*, CytHH). The electrode potential was held at E = +0.15 V for all experiments. The shaded area shows standard deviation of replicates (N =3).

The voltammogram of CycN displayed a pair of anodic and cathodic peaks (Figure 2B) consistent with Reaction (1). Based on the midway point between the two peak potentials, the formal potential (E^0′^) was calculated to be −0.009 V [51]. This E^0′^ corresponds to +0.278 V vs. NHE (normal hydrogen electrode), consistent with the reported (+0.254 V) for other cytochrome c proteins [50]. Based on these values, a potential of +0.15-0.2 V was applied to drive CycN oxidation for subsequent nicotine analysis. We further characterized the kinetics of CycN electron transfer with the electrode by varying the scan rate of CV (Figure S1). From the variation of the peak separation, we calculated the standard rate constant (k_0_) of Reaction (1) using Nicholson’s method [52]. The k_0_ was 0.7 ×10^-2^ cm s^-1^, consistent with the reported values of other cytochrome c proteins (0.35-2.4 ×10^-2^ cm s^-1^) [53, 54].

We next tested whether CycN can act as a mediator to shuttle electrons between NicA2 and an electrode. When an electrode potential of +0.15 V was applied, the rate of CycN oxidation was predicted to increase due to CycN being reduced by NicA2 in the presence of nicotine (Figure 2A):

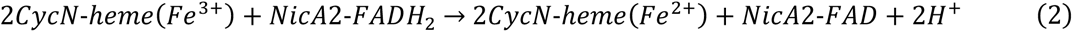

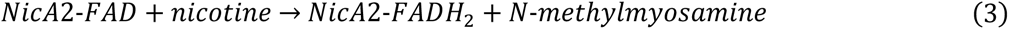

Consistent with these predicted reactions, adding nicotine to the mixture of CycN/NicA2 produced a substantial increase in anodic current (Figure 2C). In comparison, NicA2 or CycN alone showed little or no response to nicotine. We further tested if NicA2 electron transfer can be mediated by horse heart cytochrome c (CytHH) with 50.7% sequence similarity to CycN. Although CytHH displayed similar electrochemical behavior to CycN (Figure S2), the CytHH/NicA2 mixture showed no response to the addition of nicotine (Figure 2C). This result is consistent with a previous report where NicA2 reacted poorly with bovine heart cytochrome c (43% sequence similarity to CycN) [47] and with the hypothesis that CycN is adapted to NicA2 to enable efficient electron transfer.

We further determined the kinetics of NicA2 re-oxidation by CycN in the context of a sensor by estimating the rate constant of Reaction Equation (2). The addition of nicotine triggered a change in the voltammogram of CycN from a pair of redox peaks to an anodic wave characteristic of a mediator reaction [55] (Figure S3). From the limiting current of the anodic wave and treating Reaction (2) as pseudo-first-order, the rate constant was estimated to be 2.3 × 10^5^ M^-1^s^-1^ [56]. This value is consistent with the re-oxidation rate constant obtained from spectrophotometry (∼0.8 × 10^5^ M^-1^s^-1^) [47], and is comparable to that of mediators commonly used in biosensors [39].

### Highly sensitive nicotine sensor based on co-immobilization of NicA2 and CycN

We next applied CycN as a mediator to develop a second-generation nicotine sensor. NicA2 and CycN were co-immobilized onto SAM-modified gold interdigitated electrodes (IDE) by crosslinking with glutaraldehyde and chitosan. To verify sensor functionality, we conducted cyclic voltammetry at each fabrication step. Importantly, the sensor showed consistent voltammograms with those of bulk solution confirming that CycN retained redox activity and mediator capability after immobilization (Figure 3A). To maximize sensitivity, we optimized the concentrations of NicA2 (75 pmol) and CycN (1500 pmol) (Figure 3B). Protein concentrations higher than 1500 pmol did not improve sensitivity likely due to diminished diffusion of analyte (as the thickness of the protein layer increased).

**Figure 3.**
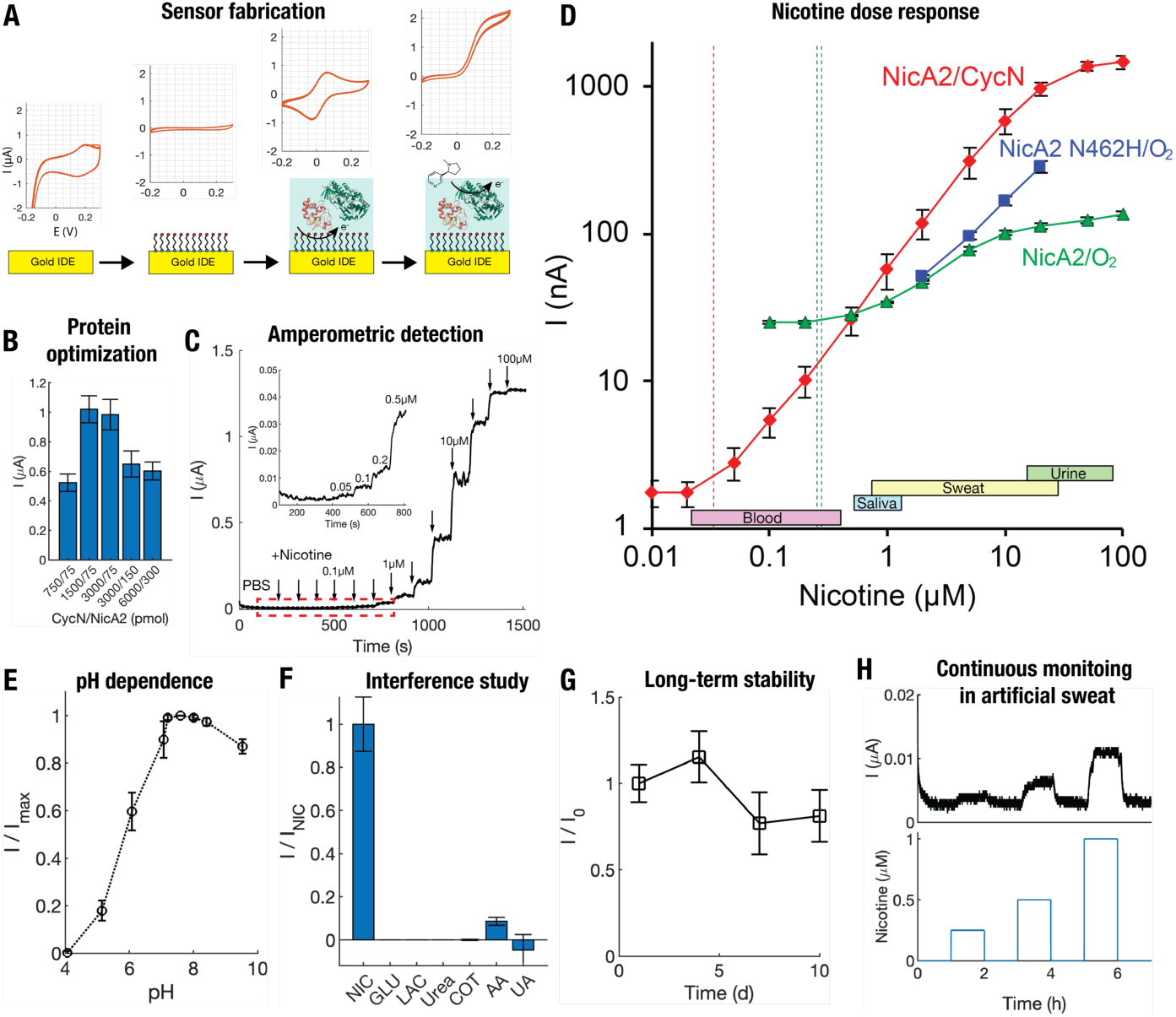
Coupling NicA2 with CycN significantly enhances sensor sensitivity. **(A)** Fabrication steps of NicA2/CycN sensor and cyclic voltammetry (CV) at each step. (From left to right) The gold interdigitated electrode (IDE) showed residue background current in PBS. Addition of a self-assembled monolayer (3-mercapto-1-propanol or 4,4′-dithiodipyridine) resulted in a decrease of capacitive current. NicA2 and CycN were next immobilized onto the electrode using chitosan as matrix and glutaraldehyde as crosslinking agent; CycN retained redox activity after immobilization as verified by CV. Finally, the sensor showed a large increase of anodic current when nicotine was added, confirming enzyme catalysis mediated by CycN. **(B)** Optimization of sensitivity by varying protein concentrations. The sensor showed highest nicotine response at 1500 and 75 pmol of CycN and NicA2, respectively. N = 3 replicates for each case. Response measured at 20 μM nicotine. **(C)** Chronoamperometry of the sensor with increasing concentrations of nicotine (0.01-100 μM) in PBS buffer. The sensor showed nicotine response down to nanomolar range (Inset). The solution was stirred throughout the experiment. The electrode potential was held at E = +0.15 V (vs screen-printed silver reference). **(D)** Nicotine dose response of the sensor and comparison with first-generation sensors based on NicA2 and its variant NicA2 N462H. Dashed lines indicate limit of detection (LOD). The sensor showed a linear range of 0.05 to 10 μM with a slope of 59.2 nA/μM. N = 3 replicates. The LOD was calculated to be 33.6 nM. The NicA2/O_2_ and NicA2 N462H/O_2_ sensors showed a sensitivity of 13.1 and 7.7 nA/μM and a calculated LOD of 0.25 and 0.28 μM, respectively. **(E)** pH-dependence of the sensor. The sensor showed maximum nicotine response at pH 7.6 and the response was stable over pH 7-9. The response decreased drastically below pH 7. N = 4 replicates. Response measured at 2 μM nicotine and normalized to pH 7.6. **(F)** Sensor performance against common biological interferents. NIC = nicotine. GLU = glucose. LAC = lactate. COT = cotinine. AA = ascorbic acid. UA = uric acid. Except for ascorbic acid (8 % of nicotine) the sensor showed little or negligible interferences from other analytes. N = 3 replicates. Response measured as nA per μM analyte and normalized to nicotine. **(G)** Stability of sensor over 10 days when stored in cold room (4 ℃) after fabrication. The sensor retained 81% of its initial response at Day 10. N = 5 replicates. Response measured at 2 μM nicotine and normalized to Day 1. **(H)** Continuous sensing in artificial sweat with physiological sweat rate. The top panel shows sensor response in a flow cell (flow rate = 2 μL/min). The bottom panel shows nicotine concentrations. The IDEs were measured in a two-electrode setup (without separate reference). E = +0.2 V.

We then extensively characterized the performance of the NicA2/CycN sensor (Figure 3C-H). The sensor showed rapid response to nicotine in a wide concentration range (10^-8^-10^-4^ M) (Figure 3C). The response was linear from 50 nM to 10 μM with a slope of 59.2 nA per μM nicotine (Figure 3D). The detection limit (LOD) was calculated to be 33.6 nM. The performance of the sensor showed significant improvement when compared with first-generation sensors based on NicA2 and an engineered NicA2 variant (N462H) with enhanced oxygen reactivity, which displayed a sensitivity of 13.1 and 7.7 nA/μM with a calculated LOD of 0.25 and 0.28 μM, respectively (Figure 3D). Thus, the LOD was improved by nearly tenfold for the NicA2/CycN sensor and two orders of magnitude compared to the first-generation sensors [41]. This was achieved by using ∼40x lower amount of enzyme, demonstrating the high efficiency of CycN. Importantly, the inclusion of CycN also enabled the sensor to operate at a much lower potential (+0.15 V) which is critical to prevent biological interferences. The nanomolar sensitivity (LOD) makes the sensor suitable for analysis of various body fluids including sweat (Figure 3D).

The sensor was further tested for its pH response, resistance to interference, and its ability for continuous monitoring. The sensor showed stable pH response between pH 7-9 (≥90% of maximum) (Figure 3E), high selectivity against potential biological interferences (<10% of nicotine) including cotinine which is the main metabolite of nicotine (Figure 3F), and long-term stability when stored at 4 ℃ (≥80% of original activity for 10 days). Finally, we demonstrated continuous monitoring of nicotine in artificial sweat using a continuous flow chamber with a flow rate of 2 μL/min, representative of physiological sweat rate [57] (Figure 3H).

### Wearable device design and sensor optimization for sweat monitoring

To deploy the nicotine biosensor for real-time on-body measurement of nicotine, we designed and fabricated a wearable device for real-time, multiplexed and wireless sweat monitoring (Figure 4A-E). The device featured a flexible microfluidic patch made of multiple skin-adhesive layers, a multiplexed sensor, and a custom printed circuit board (PCB). The microfluidic patch is designed to continuously collect and transport sweat to the sensor to enable real-time monitoring (Figure 4B-D). The skin adhesive layer of the microfluidic patch forms a flexible seal on the skin around the sweat collection area (Figure 4C, D). This configuration facilitates sweat flow into sensing chambers, driven by the positive pressure generated by sweat glands. The sensing chambers each have a volume of 3 μL. The microfluidic patch interfaces a custom multiplexed gold IDE sensor (Figure 4D). One IDE is utilized for nicotine detection in sweat, and a second IDE serves as a reference electrode to capture real-time background signals from sweat. The PCB integrates four potentiostat-on-chip integrated circuits (ICs; LMP91000) to enable multiplexed operation of up to four electrochemical sensors in parallel (Figure 4D). A Bluetooth Low Energy microcontroller unit (BLE MCU; nRF52840) is employed for data acquisition and wireless transmission (Figure 4E). The PCB is connected to the IDEs through a flexible ribbon cable and a separate connector PCB, allowing for folding and stacking of the device components to achieve a compact, low-profile design.

**Figure 4.**
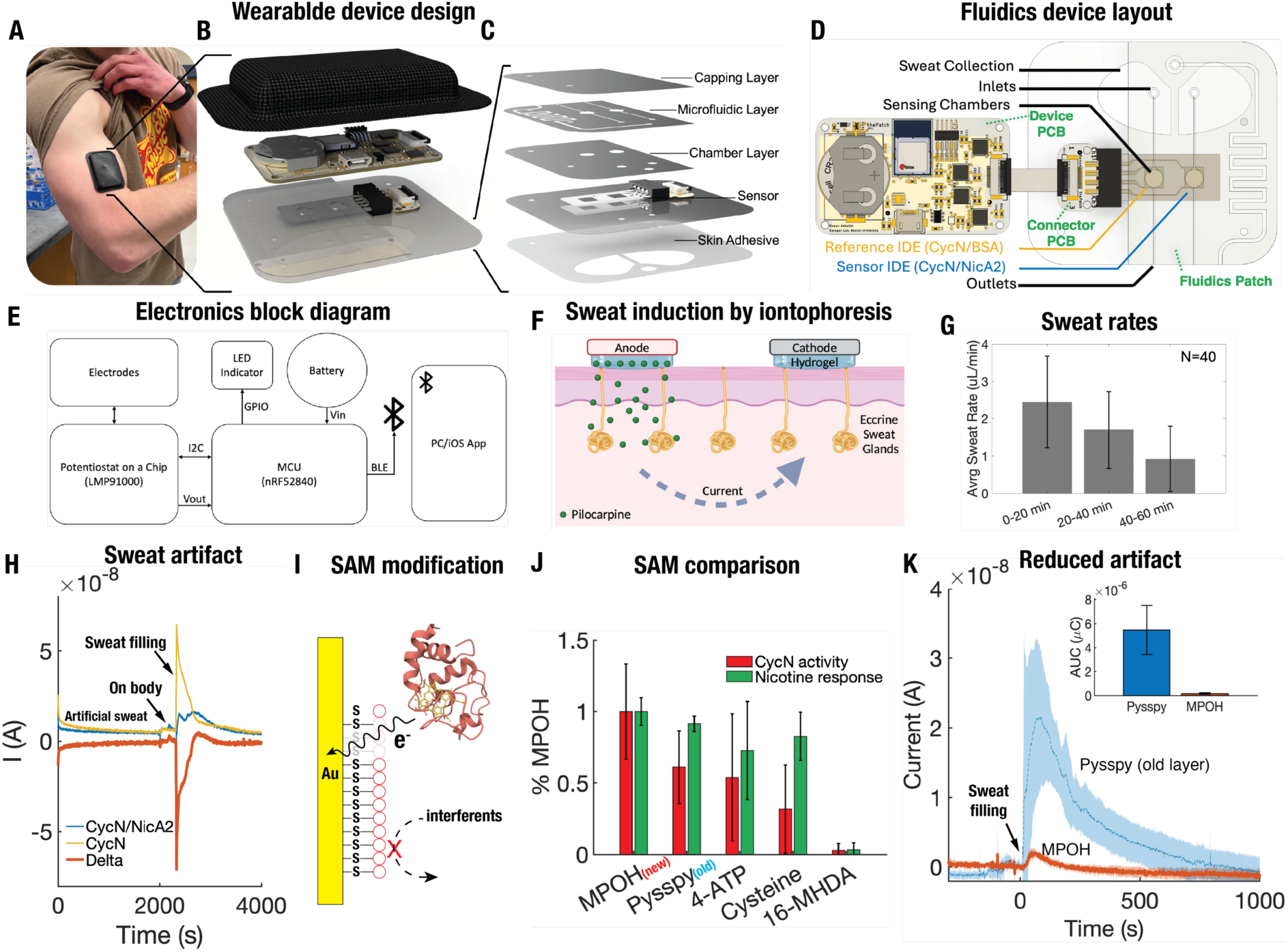
Wearable device design and sensor optimization for continuous sweat sensing on body. **(A)** Photograph of the wearable device worn on a subject’s arm. **(B)** Expanded view of the device depicting a flexible protective cover, a custom printed circuit board (PCB), and flexible microfluidic and sensor layers. **(C)** Detailed schematic of the microfluidic subsystem with labeled components. **(D)** Top view of the unfolded device, illustrating the sensor, microfluidics elements and the connection to the PCB via a flexible ribbon cable. A multiplexed gold IDE sensor was fabricated in house to provide a Reference IDE (CycN/BSA) as internal control that accounts for nonspecific signal to the Sensor IDE (CycN/NicA2). **(E)** Simplified block diagram of the electronics subsystem. **(F)** Schematic representation of sweat stimulation using iontophoresis and pilocarpine. **(G)** Average sweat rates observed at 20, 40 and 60 minutes following sweat stimulation using a commercial iontophoresis device (Macroduct). **(H)** Sensors blocked with Pysspy sensor displayed a large anodic artifact on initial channel filling on body, but not with pre-collected or artificial sweat. **(I)** Schematic illustrating the design of electrode-SAM interface for minimizing sweat interference. **(J)** Characterization of different SAM-modified electrodes for their CycN activity and nicotine response. MPOH = 3-mercapto-1-propanol. Pysspy = 4,4 ′-dithiodipyridine. 16-MHDA = 16-mercaptohexadecanoic acid. 4-ATP = 4-aminothiophenol. The standard rate constant (k_0_) of CycN and nicotine catalytic current ([NIC] = 100 μM) were normalized against MPOH. **(K**) Comparison of on-body responses of MPOH and Pysspy-blocked sensors. At t = 0 s, sweat starts filling the device. The current response was the average of 6 independent measurements (from 3 non-nicotine users). Shaded area shows standard deviations. Inset shows the area under the curve (AUC) of the artifact signal.

To passively generate sweat localized to the region of the wearable device, we utilized the cholinergic agonist pilocarpine, delivered via iontophoresis. Previous efforts to monitor sweat often induce sweat by physical exercise [58]. However, exercise may fail to produce enough sweat, introduce unnecessary participant exertion, and it is difficult to control the intensity in human participants. Local sweat stimulation using pilocarpine has been safely used as a research tool for over 30 years [59], and more recently, groups have demonstrated the use of carbachol, another cholinergic agonist, for localized and sustained sweating [58]. Pilocarpine mimics the effect of acetylcholine on eccrine sweat glands (Figure 4F), inducing sweat secretion for up to 60 minutes. Using this approach, we observed average flow rates of 2.35, 1.74, and 0.95 μL/min for the time intervals 0-20, 20-40, and 40-60 minutes following sweat stimulation, respectively (Figure 4G).

We first tested the device in non-nicotine users to determine the response of sensor to sweat without nicotine and thus establish a baseline condition for the device. However, preliminary experiments revealed a significant artifact signal when sweat first entered the sensing chambers (Figure 4H). Upon sweat filling the chambers, both the sensor (NicA2/CycN) and reference (BSA/CycN) IDEs displayed a significant transient anodic response. Notably, no such response was observed when artificial sweat or sweat collected off body was injected into the device in a continuous flow setup (Figure S4). Moreover, the magnitude and duration of the transient response varied considerably between the two IDEs. We hypothesized that the artifact signal was due to the oxidation of an electroactive species in sweat. However, testing of a range of potential candidates (urea, lactate, uric acid, ascorbic acid, pilocarpine, ethanol) failed to identify the likely species (data not shown).

To reduce interference from non-specific oxidation, one common method is to coat the sensor with a permselective membrane e.g. Nafion [60]. However, when we tested Nafion we observed a complete loss of nicotine response (Figure S5), possibly due to the acidity of the Nafion solution causing protein denaturation. An alternative approach is the modification of the electrode with a self-assembled monolayer (SAM) to act as blocking layer to prevent contact between non-specific electroactive species and the surface of the electrode [61] (Figure 4I). Our initial sensor employed 4,4′-dithiodipyridine (Pysspy) as a blocking layer. We evaluated several alternative SAMs of different chain lengths and functional groups including 3-mercapto-1-propanol (MPOH), 16-mercaptohexadecanoic acid (16-MHDA), 4-aminothiophenol (4-ATP) and cysteine. Of the SAMs tested, MPOH displayed the maximum CycN kinetics and nicotine activity (Figure 4J). Importantly, the MPOH-modified sensor also showed significantly reduced (>32 fold in AUC) sweat transient compared to the original Pysspy-blocked sensor (Figure 4K). With the MPOH-modified sensor, the average magnitude of the artifact (0.84 nA) was reduced to about one tenth of the expected nicotine-specific response (Figure 4H). We thus used MPOH for subsequent sensor development and sweat analysis.

### Real-time nicotine monitoring in sweat

We next tested the ability of the optimized nicotine sensor, deployed in the wearable device, to quantitatively measure nicotine in sweat on volunteer participants. The protocol used to test the device is shown in Figure 5A (Boston University IRB Protocol #4803E). We first stimulated sweat at the same location on the participants’ right and left forearms. The device was then applied to the site of sweat stimulation on the participants’ right forearm for one hour. In parallel, three sweat samples were collected from the site of stimulation on the left arm at 20-minute intervals for subsequent mass spectrometry analysis. Nicotine users were asked to consume nicotine as cigarettes or nicotine pouches while wearing the device. Non-users of nicotine did not consume any nicotine during the study.

**Figure 5.**
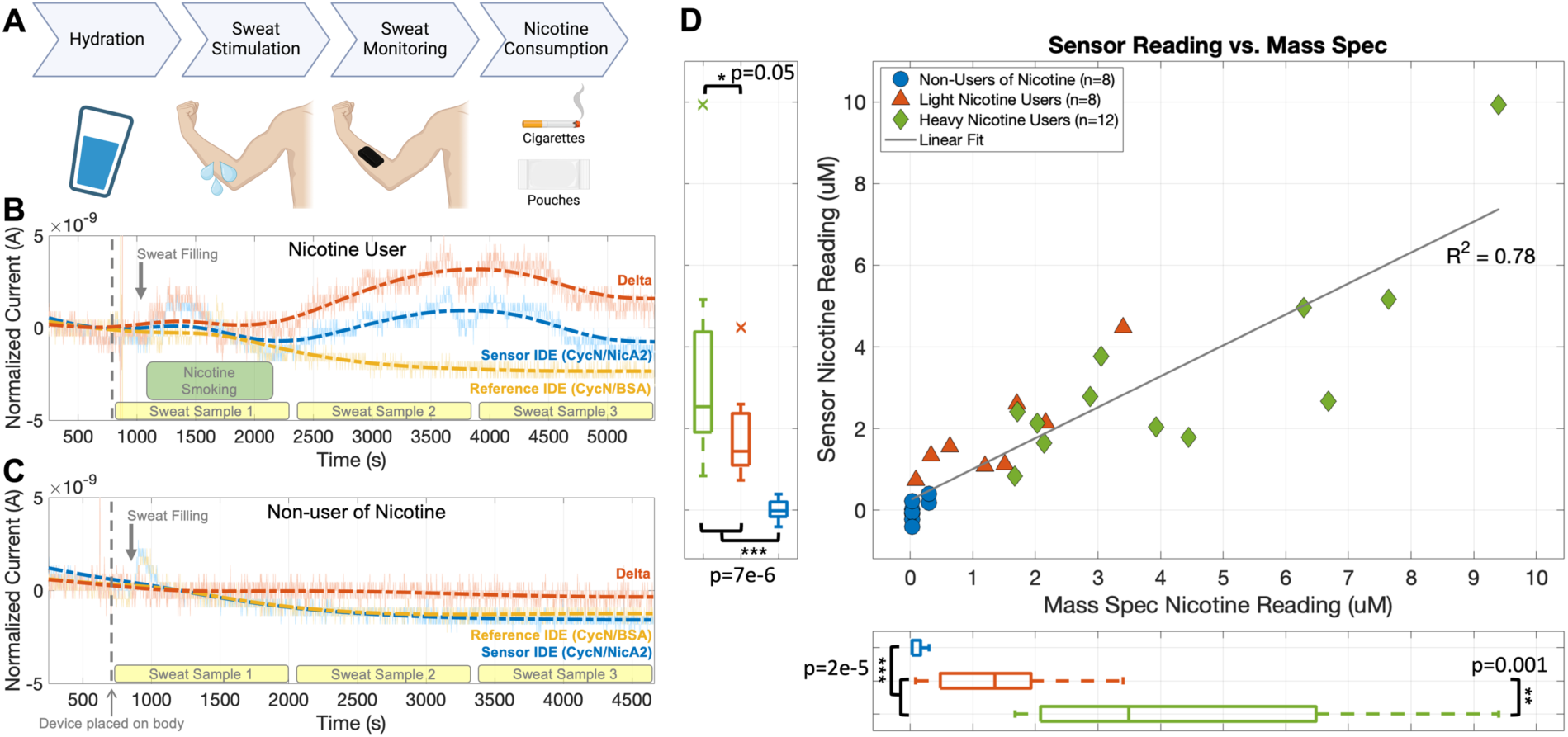
Accurate detection of nicotine on body from sweat in real-time for nicotine users and non-users. **(A)** Protocol used for testing the nicotine sensor on participants. **(B)** Real-time on-body sensor data with raw data shown as solid lines and low-pass filtered signals shown as dashed lines for a nicotine user. **(C)** Data for a nonuser. **(D)** Scatter plot comparing the mass spectrometry nicotine concentrations in collected sweat samples (x-axis) to the corresponding wearable device readings (y-axis) for all participants. Box-and-whisker plots for the three participant groups are shown below the x-axis for mass spectrometry readings and to the left of the y-axis for device readings. Asterisks indicate the significance levels of paired t-tests between the groups.

The wearable device showed a reproducible and nicotine specific signal in the sweat of nicotine users. A typical real-time device response for a nicotine user is shown in Figure 5B. The nicotine-specific signal was calculated as the difference between the nicotine (NicA2/CycN) sensor and the reference (BSA/CycN) sensor. To translate sensor signals to predictions of nicotine concentration, sensor calibration curves for each set of sensors were generated at different concentrations of nicotine and flow rates (see Methods, Figure S6). Within minutes of smoking, a nicotine signal was detected in sweat by the wearable device, the timing of which is comparable to the reported dynamics of nicotine in blood [62, 63]. For non-users of nicotine, no difference between the nicotine and reference sensors were detected, and hence no nicotine-specific signal was observed (Figure 5C).

Three populations were recruited for testing: heavy and light users of nicotine as well as non-users of nicotine. Heavy nicotine users were defined as individuals who consistently smoked 10+ cigarettes, or consumed 20+ mg of nicotine pouches, daily and smoked three or more cigarettes during the study. Light nicotine users were defined as occasional to daily users of nicotine who did not meet the criteria for heavy use, and smoked less than three cigarettes, or used a nicotine pouch, during the study. Non-users of nicotine were defined as participants who did not consume nicotine.

For each participant, we compared wearable device nicotine level measurements to mass spectrometry performed on the left-arm collected sweat samples. A total of 16 study visits were performed with 6, 5, and 5 visits of heavy, light, and non-users of nicotine, respectively. Mass spectrometry measurements for left arm sweat samples, collected over 20-minute intervals, were compared to the nicotine measurements from the wearable device averaged over the same interval. Each participant typically resulted in one or two sweat sample predictions per participant visit.

The wearable nicotine sensor measured nicotine in sweat with equal or higher analytical accuracy than mass spectrometry (Figure 5D). The sensor showed high correlation to mass spectrometry (R^2^ = 0.78) in the observed sweat nicotine levels for the three populations. The observed levels were also consistent with the reported levels of nicotine in sweat [64]. Additionally, the sensor can accurately distinguish users and non-users of nicotine. A one tailed t-test of the means of sensor nicotine readings for all users of nicotine compared to non-users showed a highly statistically significant difference with a p-value of 7e^-6^, compared to a p-value of 2e^-5^ for mass spectrometry. Moreover, heavy and light nicotine users were also differentiated by the sensor and mass spectrometry with p-values of 0.05 and 0.001, respectively. A logistic regression analysis showed perfect classification of nicotine use by the wearable device with a receiver operating characteristic (ROC) area under the curve (AUC) of 1.00, compared to 0.99 for mass spectrometry (Figure S7). In contrast, mass spectrometry misclassified one light nicotine user sample as a non-user of nicotine.

## Discussion

We report the development and deployment of a highly specific and sensitive continuous biosensor for nicotine. The biosensor comprises a microbial redox enzyme, NicA2, and its natural electron acceptor, CycN, both recovered using a genomic screen applied to the nicotine degrading bacteria *P. putida*. The biosensor uniquely pairs a redox enzyme with its natural electron acceptor and highlights the potential for improving sensor performance by using the natural electron acceptor as the mediator. Deployment of our sensor on-body with a wearable device enables real-time monitoring of nicotine in sweat. Additionally, measurements of nicotine in a study of volunteer nicotine users and non-users validates accuracy. The device demonstrates accuracy greater than or equal to gold-standard mass spectrometry for both measuring nicotine levels in sweat and differentiating nicotine users from non-users.

Our sensor represents a significant advance relative to other reported nicotine biosensors (Table S1). The sensor is highly specific to nicotine. The majority of previously reported nicotine biosensors utilize biorecognition elements or processes that are not specific to nicotine. These include sensors that respond to nicotine using enzymes whose activity is inhibited by nicotine [23–26], using nanomaterials that promote direct electro-oxidation of nicotine [28, 29], and using the nicotine-degrading capability of cytochrome P450 2B6 [20]. None of these mechanisms are specific to nicotine. Cytochrome P450 2B6, for example, degrades cotinine which is the main metabolite of nicotine [31]. By contrast, our sensor utilizes NicA2 as the biorecognition element, a redox enzyme highly specific to nicotine (binding affinity, K_m_ ∼100 nM [65]). Our sensor is also substantially more sensitive than nearly all previously reported sensors, enabling measurement over the expected concentration of nicotine in relevant biofluids (Figure 3D). Two previous sensors with comparable sensitivity lack the specificity of our sensor [28, 29]. Finally, we have validated our sensor for accurate real-time sensing on-body in an appropriately powered study of multiple volunteers compared to gold-standard mass spectrometry. Only one other study reported on-body results [20], and that report used the above cytochrome P450 2B6 based sensor. However, that work only provided four examples of on-body results without statistical validation, accuracy assessment, or comparison to gold standard measurements.

A unique aspect of our sensor is the use of the natural electron acceptor, CycN, as the mediator in a second-generation sensor design, as this has not been previously reported. An ideal mediator for an enzyme-based electrochemical sensor efficiently and selectively shuttles electrons between the redox enzyme and the electrode surface. First generation sensors use molecular oxygen to accept electrons from the enzyme but suffer from oxygen dependence and interference due to other electroactive species. Moreover, from a bioenergetics perspective oxygen is not an efficient electron acceptor and often plays the role of terminal electron acceptor in the electron transport chain [50]. The energy released by enzymatic redox reactions is typically captured by transferring released electrons to natural electron acceptors. These reducing equivalents can then be used as reducing power in respiration or biosynthesis. Direct transfer of electrons to oxygen in these cases would result in a loss of energy to the cell. Redox enzymes may thus co-evolve with a natural electron acceptor to ensure efficient, selective, and potentially, regulated electron transfer [66]. In the case of NicA2, oxygen performs poorly as co-substrate to oxidize the flavin co-factor [47]. However, CycN whose expression is co-regulated with NicA2 in response to nicotine, increases nicotine turnover by up to 10,000-fold and is proposed to be the natural *in vivo* electron acceptor for NicA2 [47]. In this report, we demonstrate that the use of CycN as a mediator for NicA2 results in an order of magnitude improvement in sensor LOD, as compared to the first-generation sensor using oxygen (Figure 3D). The results highlight the specificity of the interaction between NicA2 and CycN; an alternative cytochrome (horse heart cytochrome c) showed no electron transfer (Figure 2C). Moreover, the efficiency of CycN is reflected in a lower E^0′^ which enables the NicA2/CycN sensor to be operated at a lower applied potential relative to other published sensors (Table S1), reducing interference from non-selective redox reactions at the electrode surface.

Natural electron acceptors, in particular cytochromes c, can improve sensor performance by two means: 1) they can specifically mediate electron transfer with their enzyme partner, whereas non-specific artificial mediators may promote electron transfer with other electroactive species; 2) cytochrome c proteins can be readily co-immobilized with enzymes and thus prevent mediator leakage, a common concern for continuous monitoring and *in vivo* studies. Furthermore, the results of the analysis of NicA2 and CycN indicate that such enzyme-cytochrome pairs may be widely present in nature. This includes fusions of enzyme-cytochrome pairs that provide the possibility of engineering sensors by transferring electrons directly from the fusion protein to the electrode and thus further enhancing sensitivity. Our report provides a general strategy for improving sensor performance by the identification and use of natural electron acceptors for redox-enzymes.

Our results also underscore the value of sweat for real-time non-invasive monitoring of diverse biomarkers [2, 67]. Sweat is a promising peripheral body fluid for noninvasive biochemical monitoring [2, 67], as it is rich in biomarkers including ions, small molecules (amino acids, metabolites, hormones etc.) and proteins [57, 68]. Further, analyte concentrations in sweat have shown good correlation with their blood levels [4, 9, 69]. Sweat can be collected non-invasively and continuously, through absorption from the surface of skin [8, 70, 71]. Also, sweat can be safely and locally induced through iontophoresis to allow sampling without the need for physical exertion. Collected sweat has been analyzed off-body for both disease diagnosis and drug detection [4, 20, 72, 73].

Multiple wearable sweat monitors have been reported over the past decade and our device builds on this prior art [2–9, 20, 73] (Table S2). One important variable is the design of the microfluidics subsystem for sampling sweat. While some devices place sensors in direct contact with the skin [82–90], the use of a microfluidics subsystem limits carryover between sweat samples, minimizes potential skin toxicity of sensor components, and limits sample evaporation [72]. A dedicated microfluidics system also improves and enables monitoring of sweat sample flow to the sensor. The rate of sweat production, both natural and in response to iontophoretic stimulation, varies with time. By altering the rate of analyte flux to the sensor, sweat flowrate impacts sensor output [67]. Calibration of our nicotine sensor revealed a nearly five-fold increase in sensor output as flow rates were increased from 0.5 – 2 μL/min (Figure S6C). Accounting for this effect is crucial to sensor accuracy. Several devices have been reported that use dedicated microfluidic solutions to measure flow rate on device in real time [92–97].

Additionally, our results call attention to artifacts that can arise with real-time monitoring of sweat that are particularly relevant for compounds present at low concentration. The most striking artifact was the transient anodic signal observed from the initial sweat collected on-body (Figure 4H, K). This transient was not seen with artificial sweat or sweat collected off-body before being introduced into the device. The transient was also observed in both the sensor and non-enzyme-based reference electrode. While we hypothesize that this transient is due to a volatile electroactive substance(s) produced during initial sweating, we have not yet identified the substance(s). To our knowledge this artifact signal from sweat has not been previously reported, likely due to its scale which is small compared to sensor signals for compounds present in high concentration in sweat such as lactate, which produce microamp amperometric signals. For nicotine sensing, which generated nanoamp signals, eliminating this artifact through MPOH blocking was essential. As biosensors for other compounds present in low concentrations are developed, we anticipate a growing need to minimize nonspecific interactions while preserving mediator and sensor components [58].

We also observed substantial baseline drift, likely arising from the dynamic nature of sweat production when monitored in real-time. The magnitude of this drift was significant relative to the nicotine specific signal. It was thus necessary to correct for this artifact through real time background subtraction using a dual-sensor configuration consisting of a nicotine sensor matched with an enzyme-free reference sensor. To our knowledge, only one previous wearable sweat device implemented a similar on-body background correction [74]. As real-time biosensors for other low concentration analytes become more common, we anticipate more applications will employ similar background correction methods.

Together, our results highlight the potential of microbes as reservoirs for biorecognition elements for novel biosensors that can be deployed in range of applications including on-body for real-time sensing. In prior work, we leveraged the sensing capabilities of microbes to develop a novel progesterone sensor based on a microbial transcription factor [75–77]. Here we leverage the metabolic capabilities of microbes to develop an electrochemical sensor based on a redox enzyme and its natural electron acceptor. With advances in metagenomic screening [78, 79] and AI-powered *in silico* screening and protein engineering [80], mining the functional diversity of microbes provides one solution to the need for more diverse biosensors for wearable health devices.

## Methods

### Materials

Gold interdigitated electrodes (width/gap = 5 μm), Prussian blue-modified carbon electrode (DS710), screen-printed platinum (C550) and gold electrodes (220AT) were purchased from Metrohm DropSens (Asturias, Spain). Multiplexed gold IDEs were fabricated in the Galagan Lab. The (s)-Nicotine, 3-mercapto-1-propanol (MPOH), 4,4 ′-dithiodipyridine (Pysspy). 16-mercaptohexadecanoic acid (16-MHDA), 4-aminothiophenol (4-ATP), cysteine, chitosan, glutaraldehyde, phosphate-buffered saline, acetic acid, sulfuric acid, ethanol, dimethyl sulfoxide (DMSO), and bovine serum albumin (BSA) were purchased from various vendors. Artificial eccrine perspiration (artificial sweat) was purchased from Pickering Laboratories and was adjusted to pH 7 before use. All reagents were of analytical grade. All aqueous solutions were prepared with deionized water.

### Enzyme Expression and Purification

NicA2 WT and N462H were heterologously expressed and purified via a C-terminal poly-histidine tag in *E. coli* as previously described [81, 82]. Plasmids were transformed into *E. coli* BL21 (DE3) (New England Biolabs, MA) and cultured in lysogeny broth at 37 °C until an OD_600_ of 0.6-0.8 was reached. Cultures were induced at OD_600_ 0.6-0.8 using 1 mM isopropyl β-D-1-thiogalactopyranoside (IPTG) and the temperature was lowered to 16 °C overnight. Expression was confirmed by SDS-polysaccharide gel electrophoresis (SDS-PAGE) analysis prior to protein purification. Cell pellets were resuspended in lysis buffer [50 mM Tris (pH 8.2), 500 mM NaCl, 10 mM imidazole, and 1 mM dithiothreitol (DTT)] supplemented with 0.1 % Pierce Universal Nuclease (Thermo Scientific, MA) and one EDTA-free protease inhibitor tablet (Sigma Aldrich, MO) and lysed by microfluidization. The clarified cell lysate was passed over a HisTrap HP prepacked column (Cytiva, MA) and eluted with elution buffer [50 mM Tris (pH 8.2), 500 mM NaCl, 250 mM imidazole, and 1 mM DTT]. Purified protein was dialyzed into storage buffer [50 mM HEPES (pH 7.5), 200 mM NaCl] and flash frozen in 10% glycerol. Final FAD-bound protein products were quantified by measuring absorbance at 280 nm and 450 nm [83].

### CycN expression and purification

CycN was designed with an N-terminal signal-peptide sequence, poly-histidine tag and TEV cleavage site in pET3a and the encoding gene ordered from (Genewiz). CycN containing pET3a and pEC86, a helper plasmid encoding cytochrome maturation factors, were co-transformed into *E. coli* BL21 (DE3) (New England Biolabs, MA) and cultured in 2x yeast extract tryptone medium shaking at 220 rpm and 30 °C until an OD_600_ of 0.8-1.0 was reached. Expression was induced with 100 μM IPTG and the temperature was lowered to 17 °C overnight while shaking at 60 rpm. Expression was confirmed by SDS-PAGE and heme staining analysis [84] prior to protein purification. Cell pellets were resuspended in lysis buffer [100 μM potassium phosphate (pH 7.4), 6 μM magnesium acetate, and 20% glycerol] supplemented with 0.1 % Pierce Universal Nuclease (Thermo Scientific, MA) and one EDTA-free protease inhibitor tablet (Sigma Aldrich, MO) and lysed by sonication. The clarified cell lysate was passed over a HisTrap HP prepacked column (Cytiva, MA) and eluted by a step-gradient containing 8, 16, 40, 60 and 100% of elution buffer [40 mM HEPES (pH 7.4), 100 mM NaCl, 300mM imidazole]. Pooled fractions were dialyzed overnight with TEV protease to remove the poly-histidine tag. Cleaved CycN was separated from poly-histidine tagged proteins by loading dialyzed product onto a HisTrap HP prepacked column (Cytiva, MA) and collecting the flowthrough. Cleaved CycN was fully oxidized with 5 mM potassium ferricyanide. Oxidized CycN was loaded onto a HiPrep 16/60 Sephacryl S-300 HR column (Cytiva, MA) and eluted with storage buffer [40 mM HEPES (pH 7.4), 100 mM NaCl] and flash frozen in 10% glycerol. Final heme-bound protein products were quantified by measuring absorbance at 280 nm and 410 nm [47].

### Generation of SSN

The Enzyme Function Initiative-Enzyme Similarity Tool (EFI-EST; https://efi.igb.illinois.edu/efi-est/) was used to create an SSN for the flavin amine oxidase family using Pfam identification [85, 86]. Protein sequences (173,035) were annotated as flavin containing amine oxidoreductases by the Pfam PF01593. UniRef50 cluster ID sequences were used to compress the dataset to 23,035 sequences. An e-value of 5 was used for the initial SSN calculation. The final representative SSN was generated based on an alignment score threshold of 67 corresponding to a ∼35% identity cutoff for which an edge will connect two nodes in the SSN. Sequences were restricted to lengths of 200-1000 amino acids to omit fragments from the network. The final SSN (Figure 1A) contains 17,398 representative nodes connected by 233,907 edges. The network was visualized using the yFiles Organic Layout in Cytoscape [87].

### Generation of GNN

The Enzyme Function Initiative–Genome Neighborhood Tool (EFI-GNT; https://efi.igb.illinois.edu/efi-gnt/) was used to generate a GNN. NicA2 (UniProt ID F8G0P2) and neighboring nodes were selected from the SSN for the GNN input [85, 86]. The neighborhood size was set to 10 open reading frames preceding and proceeding the query input sequence. The minimum query-neighbor co-occurrence frequency for a spoke node was set to 20%. The resulting hub-and-spoke cluster of the GNN contained six spoke nodes (Figure 1C). The network was visualized using the yFiles Organic Layout in Cytoscape [87].

### CycN electrochemical characterization

Cyclic voltammetry was conducted using a potentiostat (µStat 4000, Metrohm DropSens) and a three-electrode setup. A gold interdigitated electrode (DropSens) modified with 3-mercapto-1-propanol was used as the working electrode. A screen-printed silver and platinum electrodes of a commercial sensor (DropSens C550) were used for reference and counter electrodes, respectively. PBS (pH 7.4) was used as background electrolyte. A CycN concentration of 100 μM was used for all experiments. For characterizing mediator capability, a NicA2 concentration of 3.3 μM and a nicotine concentration of 100 μM were used for all cases.

The formal potential (E^0’^) of the CycN redox reaction was calculated by

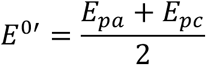

Where E_pa_ and E_pc_ are the anodic and cathodic peak potentials, respectively. The E^0’^ was converted to the NHE (normal hydrogen electrode) scale by

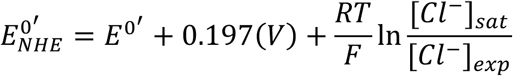

Where +0.197 V is the potential difference between saturated Ag/AgCl electrode and NHE, [Cl^-^]_sat_ and [Cl^-^]_exp_ are saturated and experimental molar concentrations of KCl (at 25 ℃), respectively.

The standard rate constant (k_0_) of the CycN redox reaction was calculated from the variation of peak separation (ΔE_p_ = E_pa_ - E_pc_) with scan rate (Figure S1) by Nicholson’s method [52].

The rate constant (k_3_) of the CycN-NicA2 reaction was calculated from the limiting current of cyclic voltammetry by

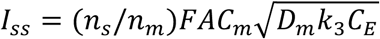

Where I_ss_ is the limiting current at E = +0.3 V, n_s_ and n_m_ are the electron number of nicotine and CycN reactions, respectively. C_m_ and C_E_ are concentrations of reduced form of CycN and NicA2, respectively. D_m_ is the diffusion coefficient of CycN and is taken from that of cytochrome c of similar size [88]. The CycN-NicA2 reaction (Reaction (2)) can be treated as a pseudo-first order reaction assuming that C_E_ remain constant in excess of nicotine [56].

### Fabrication of nicotine biosensors

The NicA2/CycN sensor was prepared on the gold interdigitated electrodes (IDE). The electrodes were cleaned with alcohol and DI and sonicated for 10-15 s. The electrodes were then preconditioned electrochemically in 0.05 M sulfuric acid (H_2_SO_4_) by cyclic voltammetry (CV, −0.5 to +1.4V) vs. a screen-printed silver reference electrode (C550, Metrohm) for 10 cycles.

For surface modification of gold electrode, the electrodes were immersed in the solution of SAM (0.001-1 M) for 30 min at room temperature depending on the SAM material (1M MPOH, 2.5 mM Pysspy, 1 mM 16-MHDA, and 3.2 mM 4-ATP dissolved in ethanol, 10 mM cysteine in DI). The electrodes were then rinsed with excess DI to remove unbound SAM. SAM solutions were usually freshly prepared for each use or refrigerated (4 °C) for short-term storage (about one week) to prevent degradation.

For electrode functionalization, a protein cocktail was freshly prepared by adding subsequently the solution of NicA2, CycN, chitosan (in 0.5 v/v% HAc) and glutaraldehyde. The cocktail was mixed thoroughly by pipetting up and down and incubated for 5-10 min before deposition. For final sensor characterization and sweat monitoring, each sensor was prepared by mixing 75 pmol NicA2 and 1500 pmol CycN. The proteins were immobilized with 0.5 μL 0.05% chitosan and 2.5 μL 0.01% glutaraldehyde. The protein cocktail was deposited onto the gold electrode and allowed to dry for 1-2 h under ambient conditions. The sensors were rinsed with excess PBS and DI to remove salts and unbound proteins. The sensors were stored in cold room (4 °C) before use.

For the NicA2/O_2_ and NicA2 N462H sensors, sensors were prepared on the platinum IDEs. Prior to enzyme deposition, equal volumes of NicA2 (3000 pmol) and 0.1 wt% chitosan and 0.05% glutaraldehyde were mixed. NicA2 was immobilized onto the IDEs by drop-casting 8-12 µL of this mixture onto the electrode. The sensors were allowed to dry 2-3 h at room temperature for solvent evaporation. Prior to use, the sensors were washed with PBS and DI and stored at 4°C.

### Microfabrication of multiplexed IDE sensor

For sweat monitoring on body, multiplexed gold interdigitated electrodes (IDEs) were fabricated using standard photolithography and lift-off techniques. Initially, the fused silica substrate underwent spin-coating of hexamethyldisilazane (HMDS) to promote surface activation and enhance resist adhesion. Subsequently, a 1.4 µm layer of AZ 5214E photoresist was spin-coated onto the substrate using a Headway Spinner. The photoresist was then patterned via exposure to UV light through a pre-fabricated mask using a Mask Aligner (MA6, i-line). A Samco PC-300 Plasma Cleaning System was employed to remove residual resist and further activate the exposed areas for improved metal adhesion.

Immediately following plasma cleaning, a 10 nm/100 nm titanium/gold (Ti/Au) layer was deposited onto the substrate via electron beam (E-beam) evaporation. The lift-off process was performed in acetone for two hours, followed by one minute of ultrasonic agitation to ensure complete removal of the resist. After confirming the electrode quality, a 10 µm thick layer of AZ4620 photoresist was applied to protect the patterned electrodes during dicing. Finally, the protective resist was removed using acetone, leaving the custom electrodes ready for use.

### Sensor characterization in vitro

Cyclic Voltammetry was conducted with a potentiostat (µStat 4000, Metrohm DropSens) in bulk solution or in a droplet of PBS buffer. A screen-printed silver and platinum electrodes (DropSens 550) were used as the reference and counter electrodes, respectively.

Chronoamperometry was performed by a potentiostat (µStat 4000, Metrohm DropSens or Emstat3, PalmSens) in bulk solution with stirring applied or in a flow cell module (Metrohm) with flow rate controlled by syringe pumps (NE1000, New Era). For characterization of NicA2/CycN sensor in the bulk solution, an electrode potential E = +0.15 V was applied (vs. screen-printed silver). For flow cell experiment, E = +0.2 V was applied across the two leads of IDE without external reference (2-electrode configuration). For nicotine dose response, current was allowed to decay to baseline for 200-300 s before adding nicotine. Data was analyzed in MATLAB and Excel.

The lower detection limit (LOD) of sensor was calculated from the intersection of linear and nonlinear portions of the nicotine dose response curve. In some cases, LOD was also estimated based on the equation:

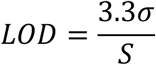

Where *σ* is the standard deviation of blank (no nicotine), S is the slope of the linear portion of the nicotine-dose response curve.

### Wearable Device Design and Fabrication

The electronics subsystem of the wearable device was developed in-house (Figure S8). It employs four electrochemical analog front-end (AFE) integrated circuits (ICs) (LMP91000, Texas Instruments) for multiplexed electrochemical sensing. A Bluetooth Low Energy (BLE) microcontroller unit (MCU) (nRF52840, Nordic Semiconductors) facilitates low-power data acquisition and wireless transmission. Printed circuit boards (PCBs) were designed in-house, fabricated by PCBWay, and soldered in-house using a reflow oven. Components for the PCBs were sourced from Mouser Electronics and DigiKey (Table S3). Backend software was developed in C++ using the PlatformIO Arduino framework and the Nordic nRF52 platform, while frontend software was implemented using MATLAB.

The disposable, flexible microfluidics patch was fabricated by laser-cutting layers of skin-compatible adhesive (ARcare® 90445Q, Adhesives Research) and polystyrene (PS) sheets to create spacing. The low thickness of the adhesive and PS layers resulted in a sensing chamber volume of 3.6 µL, enabling detection with minimal sweat samples.

### Wearable Device Testing Protocol

Light and heavy nicotine users, as well as non-users of nicotine, volunteered to participate in this study. The protocol was reviewed and approved by the Boston University Institutional Review Board (IRB) (IRB Protocol #4803E). Heavy nicotine users were defined as individuals who consistently smoked ≥ 10 cigarettes or consumed ≥ 20 mg nicotine pouches daily and smoked ≥ 3 cigarettes during the study. Light nicotine users were defined as occasional or daily users of nicotine who did not meet the criteria for heavy use and smoked 1-2 cigarettes or used a nicotine pouch during the study. Non-users of nicotine were defined as participants who do not consume any nicotine in the past and during the study.

Sweat was locally stimulated using an FDA-approved iontophoresis device (Macroduct, ELITech Group). The Macroduct Advanced Sweat Collection System uses iontophoresis to deliver a small dose (<= 1.1 mg, per the manufacturer) of pilocarpine over a 5-minute period. After sweat stimulation, the wearable device was applied to the participant’s forearm at the location of stimulation and started on-body measurement. At the same time, sweat was stimulated and samples were collected separately from the same location on a second arm for mass spectrometry analysis. Typically, a total of 3 sweat samples were collected for each participant at intervals of 20 min after sweat stimulation. Sweat samples that had an average flow rate below 0.5 μL/minute were excluded from nicotine prediction, due to the limited sensor response at very low flow rates. Additionally, the first sweat sample for all participants was not included since sweat filling occurs 5-10 minutes following device application; hence the sensor is unable to provide a nicotine reading prior to sweat filling.

Nicotine users were given the option to either smoke cigarettes (Marlboro Gold) or consume nicotine pouches (ZYN, Chill or Smooth flavor, 3 or 6 mg). Due to the slower absorption of nicotine from pouches [62], participants who chose nicotine pouches were instructed to place a pouch in their mouth 25–30 minutes before the wearable device was applied and to keep it in for one hour. Participants who chose to smoke cigarettes were instructed to begin smoking 7–15 minutes after the device was applied, with the option to smoke one or more cigarettes at their discretion. Non-users of nicotine did not intake any nicotine and remained sedentary throughout the data collection period.

The nicotine sensor was preconditioned to establish a baseline by filling device with artificial sweat prior to applying the device on body. (The presence or absence of this step did not impact the presence of the anodic transient described in the main text). A bias voltage of E = +0.2 V was applied between the two leads of the IDE for both the sensing (NicA2/CycN) and reference (BSA/CycN) IDEs. Following the device’s application, nicotine was continuously monitored for ∼1 h. Meanwhile, sweat pH was determined by loading a pH indicator dye (Bromocresol Purple) from a separate channel next to the sensing channels (in most cases pH was found to be between pH 7-8). After on-body measurement, the sensor was calibrated in a flow cell with artificial sweat to calculate sweat nicotine concentration (Figure S6). Based on sweat rate estimation, the sensor was calibrated at a flow rate of 1 and 0.5 μL/min for sweat samples collected at 20-40 min and 40-60 min, respectively. From the calibration curves, nicotine was calculated by averaging the differential (‘Delta’) and baseline-subtracted signal from the same time intervals as sweat samples.

### LC–MS analysis of sweat samples

Sweat samples were diluted 10x in 0.1% formic acid and anonymized prior to analysis. A portion of collected sweat from non-users of nicotine was pooled and spiked with nicotine to generate the nicotine standards. Analysis of sweat samples was performed on a Hybrid Triple Quadrupole/Linear ion trap mass spectrometer, SCIEX 4000 QTRAP. The samples were run in positive mode with a reverse-phase C18 Zorbax Eclipse (2.1 x 50 mm, Agilent). Separation was achieved by a flow rate of 0.2 mL/min and an isocratic mobile phase gradient for 3 minutes. The monitored transition was 163 → 132 m/z with a collision energy of 20 eV.

## Acknowledgements

The authors would like to acknowledge the BU Biomedical Engineering Core Facilities (BECF), the BU Chemical Instrumentation Center (CIC), the BU Optoelectronic Processing Facility (OPF), and the BU Center for Nanoscale Systems (CNS). Z.Z., A.A., H.Z. U.K., R.C., and J.E.G received support from DOD/ONR (Awards N000142112433, N000142212689); A.A., H.Z., and J.E.G. received support from DASA (Contract No. DSTL0000016713); U.K., R.C., J.E.G. received support from NSF (Award 2126992).

## Author contributions

(Based on https://www.niso.org/publications/z39104-2022-credit)

Conceptualization: Z.Z., A.A., C.K., K.N.A., M.W.G., J.E.G.

Data curation: Z.Z., A.A., K.H.,

Formal analysis: Z.Z., A.A., K.H., J.E.G.

Funding acquisition: C.K., K.N.A., M.W.G., J.E.G.

Investigation: Z.Z., A.A., K.H., H.Z., U.K., R.C.

Methodology: Z.Z., A.A., K.H., J.E.G.

Project administration: Z.Z., A.A., C.K., K.N.A., M.W.G., J.E.G.

Resources: A.A., K.H., H.Z., M.D.C.P., L.L.

Software: Z.Z., A.A., J.E.G.

Supervision: C.K., K.N.A., M.W.G., J.E.G.

Visualization: Z.Z., A.A., K.H., J.E.G.

Writing: Z.Z., A.A., K.H., K.N.A., M.W.G., J.E.G.

Reviewing and Editing: Z.Z., A.A., C.K., K.N.A., M.W.G., J.E.G.

## Supplementary information

**Table S1.** Comparison of the nicotine biosensor with reported nicotine sensors.

| biorecognition | specific to nicotine? | type | applied potential (vs. Ag/AgCl) | LOD ($\mu\text{M}$) | detection range | response time | application/sample | ref |
| --- | --- | --- | --- | --- | --- | --- | --- | --- |
| NicA2/CycN | yes | amperometric | +0.15 V | 0.036 | $10^{-8}$ to $10^{-4}$ M | 10 s | Sweat on body | This work |
| NicA2/O<sub>2</sub> | yes | amperometric | -0.2 V | 3.36 | $10^{-6}$ to $10^{-4}$ M | not determined | Artificial sweat/urine | [41] |
| Nicotine tetra(chloro)phenylborate | yes | potentiometric | not applicable | 10 | $10^{-5}$ to 0.08 M | 30 s to 2 min | tobacco products | [21] |
| Nicotine 5-nitrobarbiturate | yes | potentiometric | not applicable | 10 | $10^{-5}$ to $10^{-2}$ M | 30 s | tobacco products | [22] |
| Choline oxidase with 1,4-benzoquinone as mediator | no | amperometric | +0.5 V | 10 | $10^{-5}$ to $10^{-3}$ M | 10 s | tobacco samples | [24] |
| Butyrylcholinesterase/choline oxidase | no | amperometric | +0.6 V | 1 | $10^{-6}$ to $10^{-4}$ M | 2 min | tobacco and pharmaceutical samples | [25] |
| Acetylcholinesterase/choline oxidase | no | amperometric | not specified | 60 | $10^{-5}$ to $10^{-3}$ M | 5 min | phosphate buffer | [26] |
| Cytochrome P450 2B6 | no | amperometric | +0.8 V | 16 | 0 to 20 $\mu\text{M}$ | not specified | Sweat on body | [20] |
| Polydopamine functionalized reduced graphene oxide-gold nanoparticle nanocomposite | no | amperometric | +0.93 V | 0.015 | 0.05 to 500 $\mu\text{M}$ | 7 s | tobacco products | [28] |
| Nitrogen-doped graphene sheets | no | amperometric | +0.85 V | 0.047 | 0 to 500 $\mu\text{M}$ | not specified | tobacco products | [29] |
| Acetylcholinesterase | no | potentiometric | not applicable | 1000 | $10^{-3}$ to $10^{-1}$ M | 4 min | buffer | [23] |
| Boron-doped diamond | no | voltammetric (square wave) | +0.8 to +1.5V | 30 | $10^{-6}$ to $10^{-4}$ M | not specified | tobacco products | [27] |

**Table S2.** Comparison of published fully integrated sweat wearable biosensors.

| Year | Integrates Sweat Background Sensor? | Sweat Fluidics vs. Direct Skin Interface | Chamber Volume | Fluidics Material | Device Adherence Method | Capable of Continuous Sensing? | Analyte/s Sensed | Sweating Source | Reported Human Testing Sample Size | Device Reading Compared to Gold Standard? | Research Group | Ref |
| --- | --- | --- | --- | --- | --- | --- | --- | --- | --- | --- | --- | --- |
| 2014 | No | Direct | N/A | N/A | Arm Strap | Yes | Na<sup>+</sup> | Cycling | 2 | No | Joseph Wang | [89] |
| 2016 | No | Direct | N/A | N/A | Wrist/Headband | Yes | Glu/Lac/K<sup>+</sup>/Na<sup>+</sup> | Cycling/Running | 6 | Partially | Ali Javey | [90] |
| 2016 | No | Fluidics | 3.5uL | PMMA (Rigid) | Arm Strap | No (limited to 344uL volume) | Na<sup>+</sup> | Cycling | 1 | No | Dermot Diamond | [91] |
| 2016 | No | Direct | N/A | N/A | Adhesive | Yes | Lac | Cycling | 3 | No | Joseph Wang & Patrick Mercier | [92] |
| 2016 | No | Direct | N/A | N/A | Adhesive | Yes | Alcohol | Local Stimulation (On Device) | 2 | No | Joseph Wang | [93] |
| 2016 | No | Direct | N/A | N/A | Arm/Headband | Yes | Ca<sup>2+</sup>/pH | Cycling | 1 | Yes | Ali Javey | [94] |
| 2017 | No | Direct | N/A | N/A | Wristband | Yes | Na<sup>+</sup>/Cl<sup>-</sup> | Local Stimulation (On Device) | 9 | No | Ali Javey & Ronald W. Davis | [95] |
| 2017 | No | Fluidics | ~8uL | PDMS/Adhesive (Flexible) | PDMS Bandage/Adhesive | Yes | Glu/Lac | Cycling | 2 | No | Joseph Wang | [96] |
| 2018 | No | Direct | N/A | N/A | Adhesive | Yes | Glu/Alcohol | Local Stimulation (On Device) | 2 | No | Joseph Wang | [97] |
| 2018 | No | Fluidics | ~8uL | PDMS/PET (Flexible) | PDMS Wristband | Yes | pH/K<sup>+</sup>/Cl<sup>-</sup>/Na<sup>+</sup> | Cycling | 3 | No | Ali Javey | [98] |
| 2018 | No | Fluidics | ~8uL | PDMS/Adhesive (Flexible) | PDMS Bandage/Adhesive | Yes | Na<sup>+</sup>/K<sup>+</sup> | Cycling | 3 | No | Joseph Wang | [99] |
| 2018 | No | Direct | N/A | N/A | PDMS Wristband | Yes | Caffeine | Cycling | 1 | No | Ali Javey | [100] |
| 2020 | No | Direct | N/A | N/A | Arm Band | Yes | Nicotine | Cycling | 4 | No | Ali Javey | [101] |
| 2020 | No | Fluidics | 2uL | Adhesive/Polyimide/PET (Flexible) | Adhesive | Yes | Uric Acid and Tyrosine | Cycling | 25 | Yes | Wei Gao | [102] |
| 2020 | No | Fluidics | 4uL | Adhesive/PET (Flexible) | Adhesive | Yes | Glu/Lac | Local Stimulation (On Device) | 2 | No | Sam Emaminejad | [103] |
| 2022 | Yes | Fluidics | Not Reported | Adhesive/PET (Flexible) | Adhesive | Yes | Cortisol/pH | Local Stimulation (External Device) | 1 | No | Sam Emaminejad | [74] |
| 2022 | No | Fluidics | 2.36uL | Adhesive/PET (Flexible) | Adhesive | Yes | Na<sup>+</sup>/Amino Acids | Local Stimulation (On Device) | 49 | No | Wei Gao | [104] |
| 2023 | No | Fluidics | Not Reported | Adhesive/PET (Flexible) | Adhesive | Yes | C-Reactive Proteins/pH | Local Stimulation (On Device) | 4 | No | Wei Gao | [105] |
| 2024 | No | Fluidics | 2.06uL | Adhesive/PET/PDMS (Flexible) | Adhesive | Yes | Lac/UA/Glu/Na<sup>+</sup>/K<sup>+</sup>/NH<sub>4</sub><sup>+</sup> | Local Stimulation (On Device) | 10 | No | Wei Gao | [106] |
| 2025 | Yes | Fluidics | 3uL | Adhesive/PS Sheet (Flexible) | Adhesive | Yes | Nicotine | Local Stimulation (External Device) | 16 | Yes | James Galagan | This work |

**Table S3.** Bill of Materials for the Device PCB.

| Part | Value | Device | Package | Description |
| --- | --- | --- | --- | --- |
| B1 | BK-912-TR | BK-912-TR | BK-912-TR_MPD | Coin cell battery holder 20mm |
| C1 | 0.1uF | C-US_3DC0402 | C0402 | CAPACITOR, American symbol |
| C2 | 0.1uF | C-US_3DC0402 | C0402 | CAPACITOR, American symbol |
| C3 | 10nF | C-US_3DC0402 | C0402 | CAPACITOR, American symbol |
| C4 | 0.1uF | C-US_3DC0402 | C0402 | CAPACITOR, American symbol |
| C5 | 4.7uF | C-US_3DC0402 | C0402 | CAPACITOR, American symbol |
| C6 | 0.1uF | C-US_3DC0402 | C0402 | CAPACITOR, American symbol |
| C7 | 0.1uF | C-US_3DC0402 | C0402 | CAPACITOR, American symbol |
| C8 | 0.1uF | C-US_3DC0402 | C0402 | CAPACITOR, American symbol |
| C9 | 0.1uF | C-US_3DC0402 | C0402 | CAPACITOR, American symbol |
| C10 | 1uF | C-USC0402 | C0402 | CAPACITOR, American symbol |
| C11 | 1uF | C-USC0402 | C0402 | CAPACITOR, American symbol |
| C12 | 1uF | C-USC0402 | C0402 | CAPACITOR, American symbol |
| C13 | 1uF | C-USC0402 | C0402 | CAPACITOR, American symbol |
| CR1 | MM3Z3V3B | MM3Z3V3B | SOD-323FL_477AB_ONS |  |
| CU1 | 15nF | C-US_3DC0402 | C0402 | CAPACITOR, American symbol |
| CU2 | 15nF | C-US_3DC0402 | C0402 | CAPACITOR, American symbol |
| CU3 | 15nF | C-US_3DC0402 | C0402 | CAPACITOR, American symbol |
| CU4 | 15nF | C-US_3DC0402 | C0402 | CAPACITOR, American symbol |
| D1 | APHF1608LSEEQBDZGKC | APHF1608LSEEQBDZGKC | LED_APHF1608LSEEQBDZGKC | Led Rgb 0603 Smd |
| J1 | ZX62D-B-5PA8(30) | ZX62D-B-5PA8(30) | CONN_ZX62D-B-5PA8(30)_HIR |  |
| J2 | 6.8711E+10 | 6.8711E+10 | CONN_68711214522_WRE |  |
| J3 | FTSH-105-01-L-DH | FTSH-105-01-L-DH | SAMTEC_FTSH-105-01-L-DH |  |
| R1 | 10K | R-US_3DR0402 | R0402 | RESISTOR, American symbol |
| R2 | 100k | R-US_3DR0402 | R0402 | RESISTOR, American symbol |
| R3 | 100k | R-US_3DR0402 | R0402 | RESISTOR, American symbol |
| R4 | 160 | R-US_R0402 | R0402 | RESISTOR, American symbol |
| R5 | 160 | R-US_R0402 | R0402 | RESISTOR, American symbol |
| R6 | 160 | R-US_R0402 | R0402 | RESISTOR, American symbol |
| R7 | 160 | R-US_R0402 | R0402 | RESISTOR, American symbol |
| RB | 325 | R-US_3DR0402 | R0402 | RESISTOR, American symbol |
| RG | 325 | R-US_3DR0402 | R0402 | RESISTOR, American symbol |
| RR | 750 | R-US_3DR0402 | R0402 | RESISTOR, American symbol |
| RU1 | 2.2M | R-US_3DR0402 | R0402 | RESISTOR, American symbol |
| RU2 | 2.2M | R-US_3DR0402 | R0402 | RESISTOR, American symbol |
| RU3 | 2.2M | R-US_3DR0402 | R0402 | RESISTOR, American symbol |
| RU4 | 2.2M | R-US_3DR0402 | R0402 | RESISTOR, American symbol |
| U1 | LMP91000SD/NOPB | LMP91000SD/NOPB | SDA14B |  |
| U2 | LMP91000SD/NOPB | LMP91000SD/NOPB | SDA14B |  |
| U3 | LMP91000SD/NOPB | LMP91000SD/NOPB | SDA14B |  |
| U4 | LMP91000SD/NOPB | LMP91000SD/NOPB | SDA14B |  |
| U5 | BMD-340-A-R | BMD-340-A-R | BMD-340-A-R_RIG |  |
| U6 | MAX3203EETT+T | MAX3203EETT+T | 21-0137_T633+2_MXM |  |
| U7 | LDS3985M33R | LDS3985M33R | SOIC_85M33R_STM |  |

**Figure S1.**
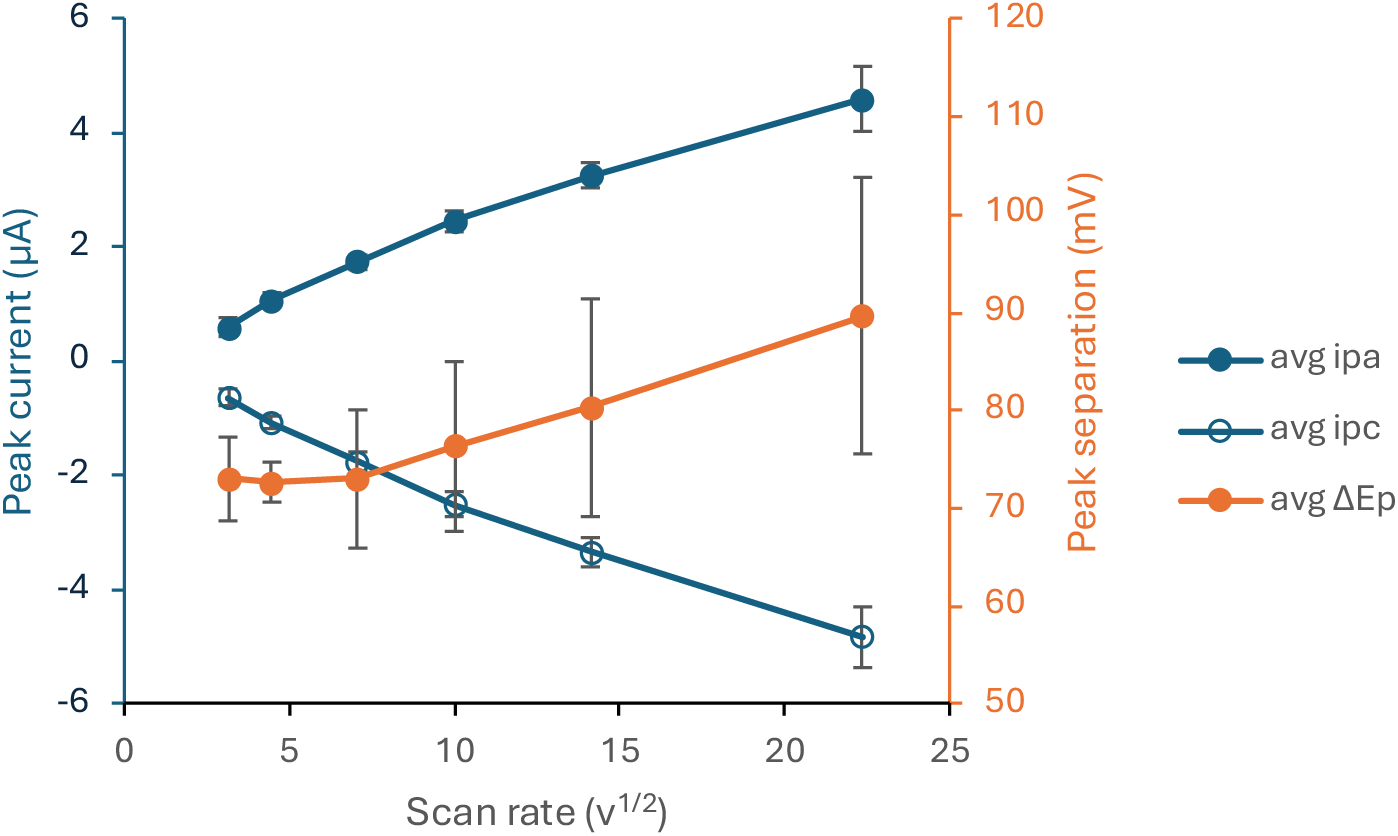
Characterization of CycN kinetics using cyclic voltammetry (CV). Showing dependence of peak currents and peak separation on scan rate. Ipa = anodic peak current. Ipc = cathodic peak current. ΔEp = peak separation. Scan rate is in mV/s. CV is conducted using PBS as background electrolyte with 100 μM CycN. Gold interdigitated electrode was modified with 3-mercapto-1-propanol. N = 3 electrodes.

**Figure S2.**
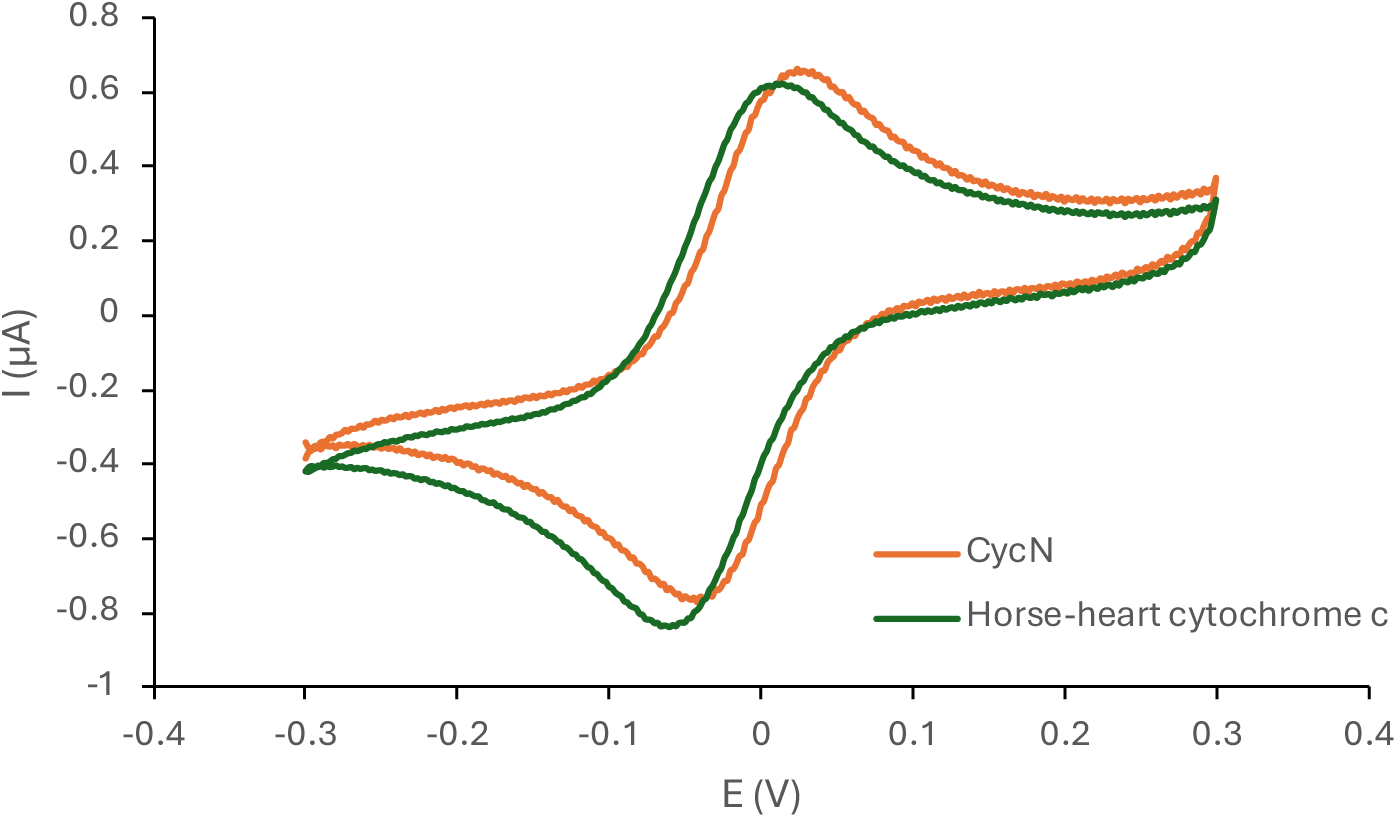
Compare CycN and another horse-heart cytochrome c (CytHH) using CV. Both displayed reversible oxidation/reduction peaks with similar standard reduction potentials around 0 V. CV is conducted using PBS as background electrolyte with 100 μM CycN or CyctHH. Gold interdigitated electrode was modified with 3-mercapto-1-propanol. A screen-printed silver electrode (Dropsens C550) was used as reference electrode. Scan rate = 50 mV/s.

**Figure S3.**
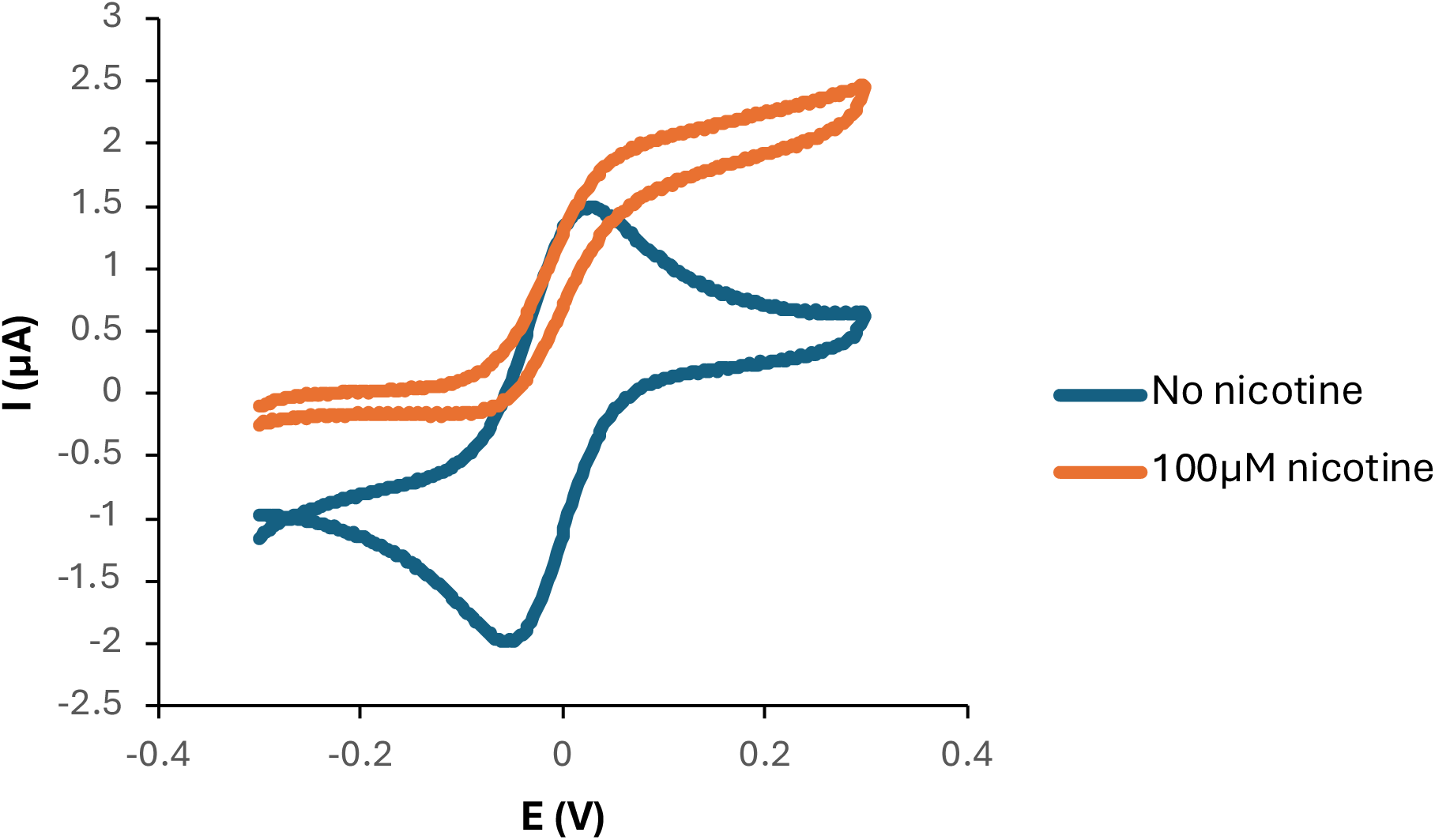
Mediation of NicA2 electrocatalysis by CycN. Upon adding nicotine to the NicA2/CycN mixture, the voltammogram transitioned from a pair of redox peaks to an anodic wave indicative of a mediator reaction. From the limiting current (Iss at E = +0.3 V), the re-oxidation rate constant (k_3_) can be estimated. A screen-printed silver electrode (Dropsens C550) was used as reference electrode. Scan rate = 50 mV/s. PBS buffer was the electrolyte. [CycN] = 100 μM. [NicA2] = 3.3 μM. Note that nicotine was provided in excess amount ([nicotine] >> K_m_) to saturate enzyme.

**Figure S4.**
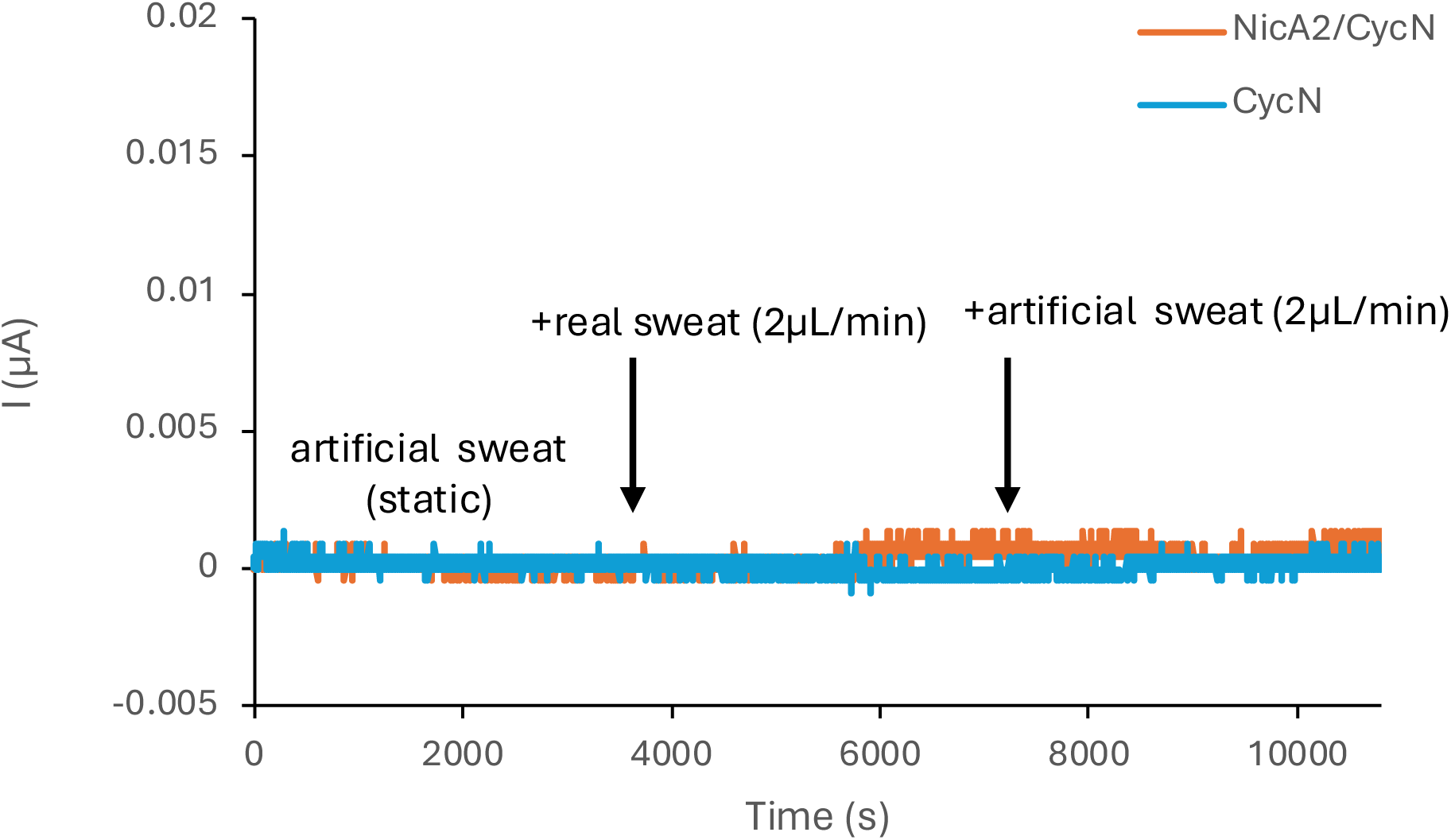
Amperometric response of multiplexed IDE sensor in a flow cell. E = +0.2 V (2-electrode setup). The sensor was initially preconditioned in artificial sweat (pH 7). At t = 3600 s, real sweat collected off body by exercise was injected into the sensor at a flow rate of 2 μL/min. At t = 7200 s, artificial sweat was again injected into the sensor at a flow rate of 2 μL/min.

**Figure S5.**
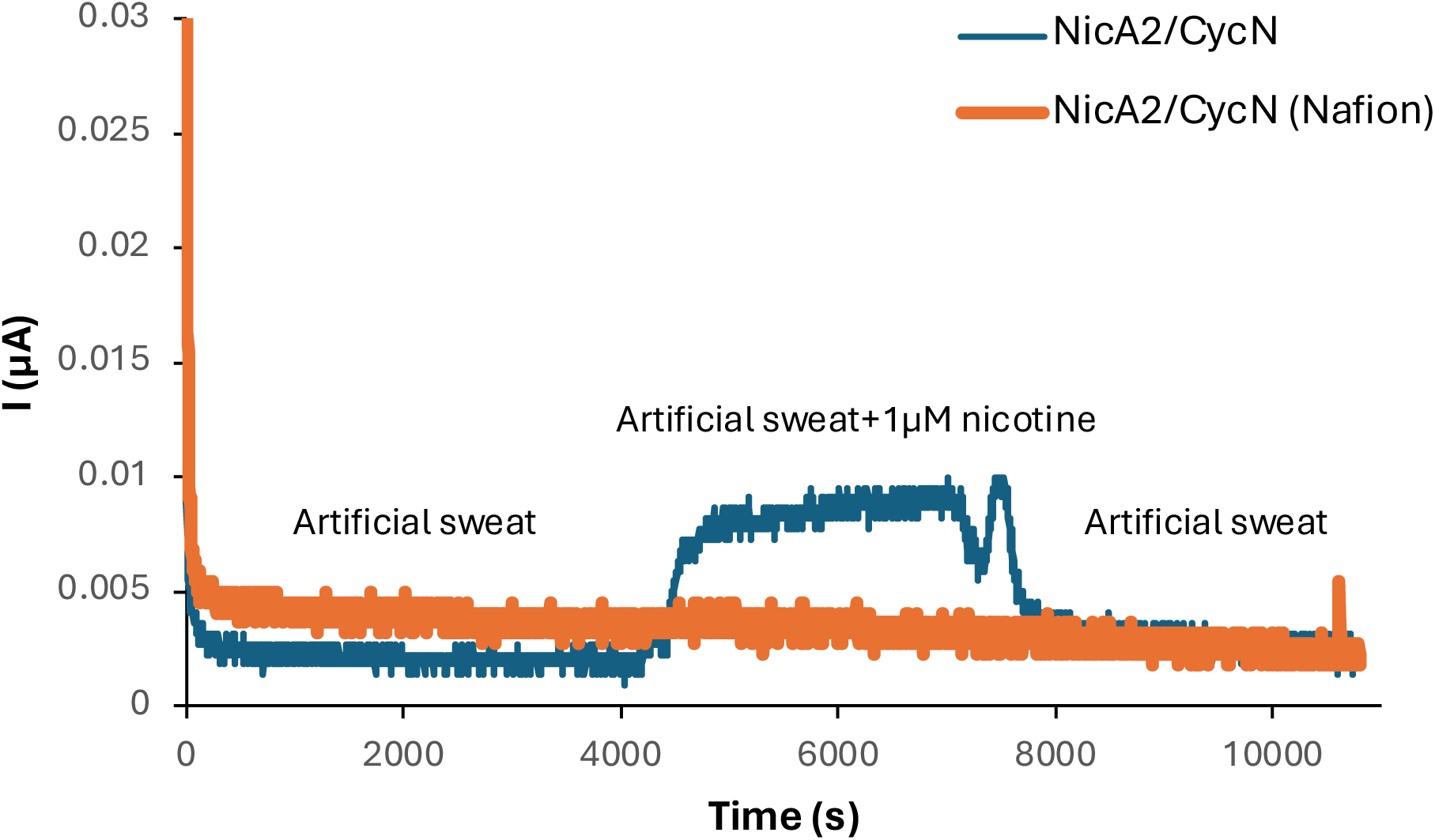
Effect of Nafion coating on the nicotine sensor. E = +0.2 V (2-electrode setup). The sensor was initially preconditioned in artificial sweat (pH 7). At t = 3600 s, nicotine-spiked sweat was injected into the sensor at a flow rate of 2 μL/min. At t = 7200 s, artificial sweat was again injected into the sensor at a flow rate of 2 μL/min.

**Figure S6.**
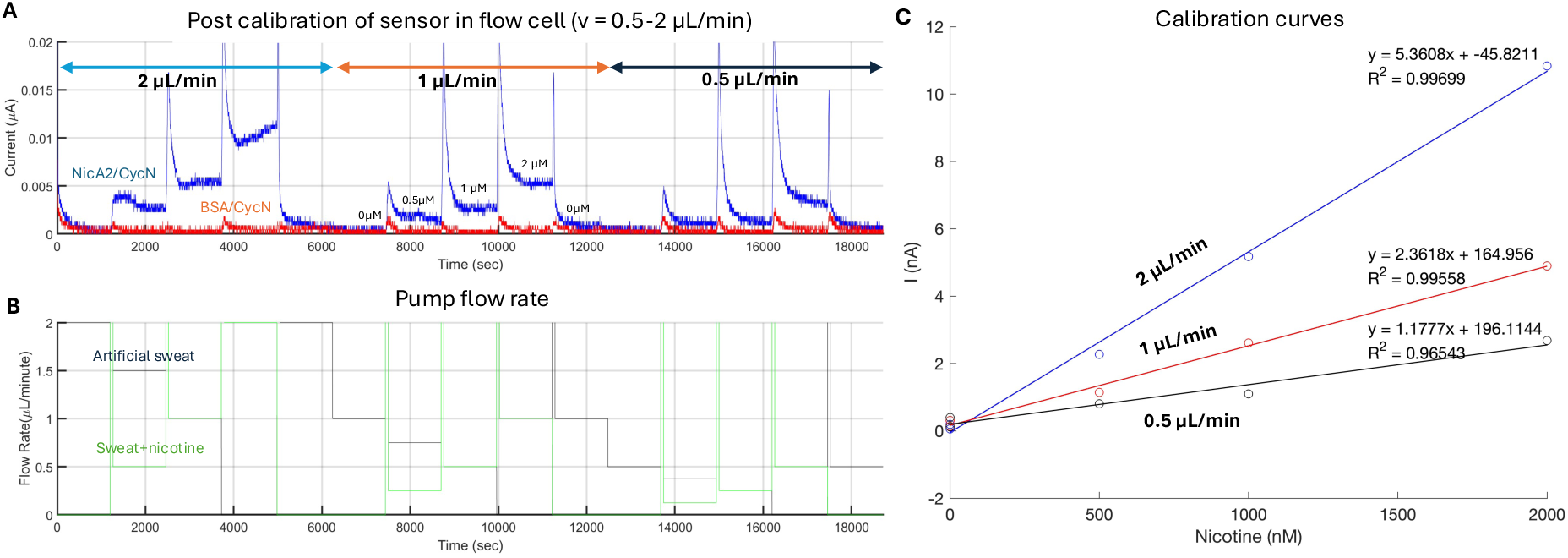
Post calibration of the nicotine sensor after on-body measurement. **(A)** The sensor was calibrated in artificial sweat (pH 7) with varying nicotine concentrations (0-2 μM) and flow rates (0.5-2 μL/min). **(B)** Flow rates of individual pump used to set nicotine levels. **(C)** Calibration curves of sensor at different flow rates. Each point was calculated by averaging the differential signal (‘Delta’) between the sensing (NicA2/CycN) and reference (BSA/CycN) electrodes in the last minute of each nicotine concentration step.

**Figure S7.**
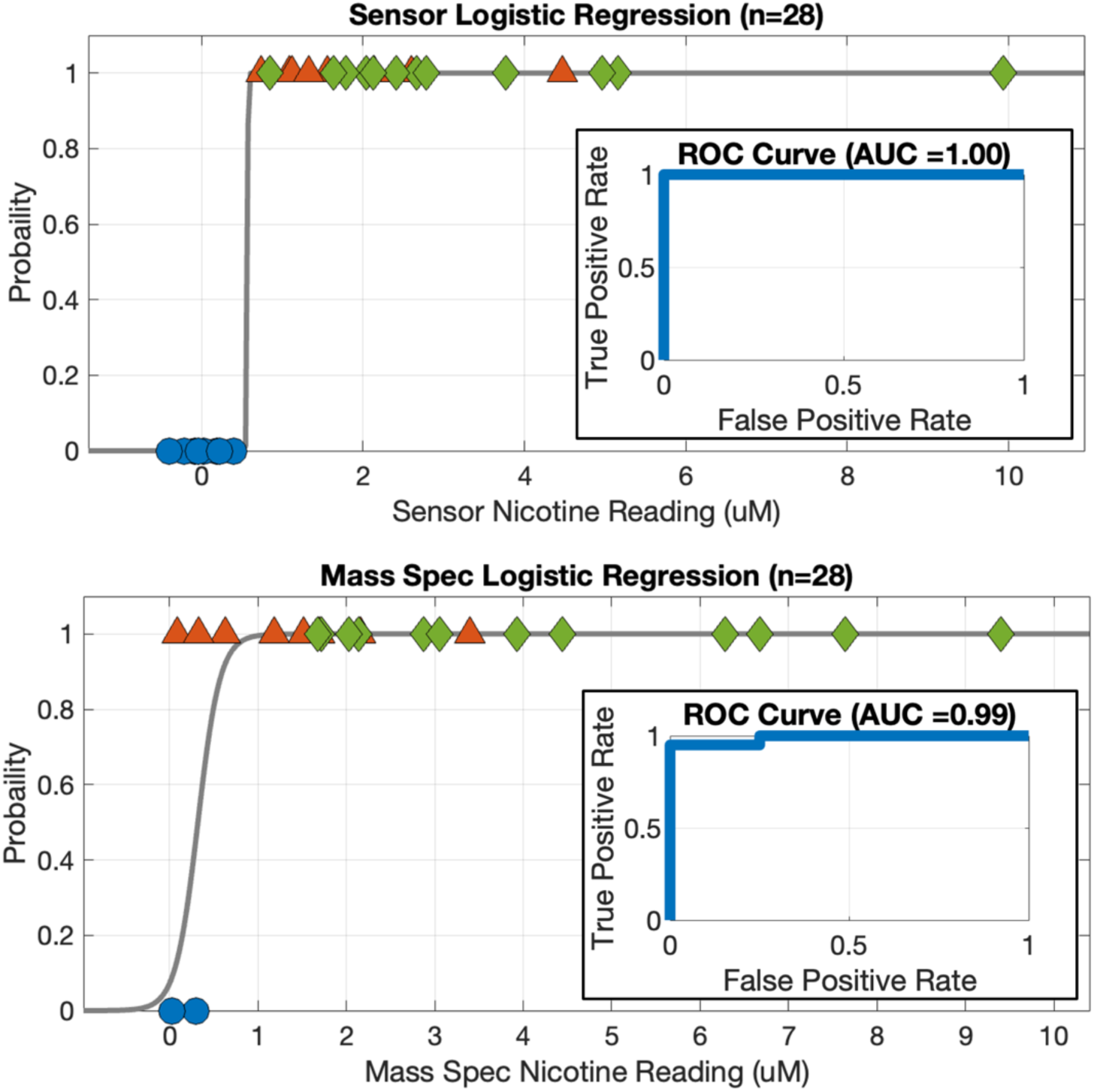
Logistic regression and receiver operator characteristic (ROC) analyses of our wearable device sensor and mass spectrometry in predicting the probability of a participant being a nicotine user. Our device displayed a perfect area under the curve (AUC = 1) characteristic, indicating that our device has been highly accurate in predicting nicotine users from nonusers.

**Figure S8.**
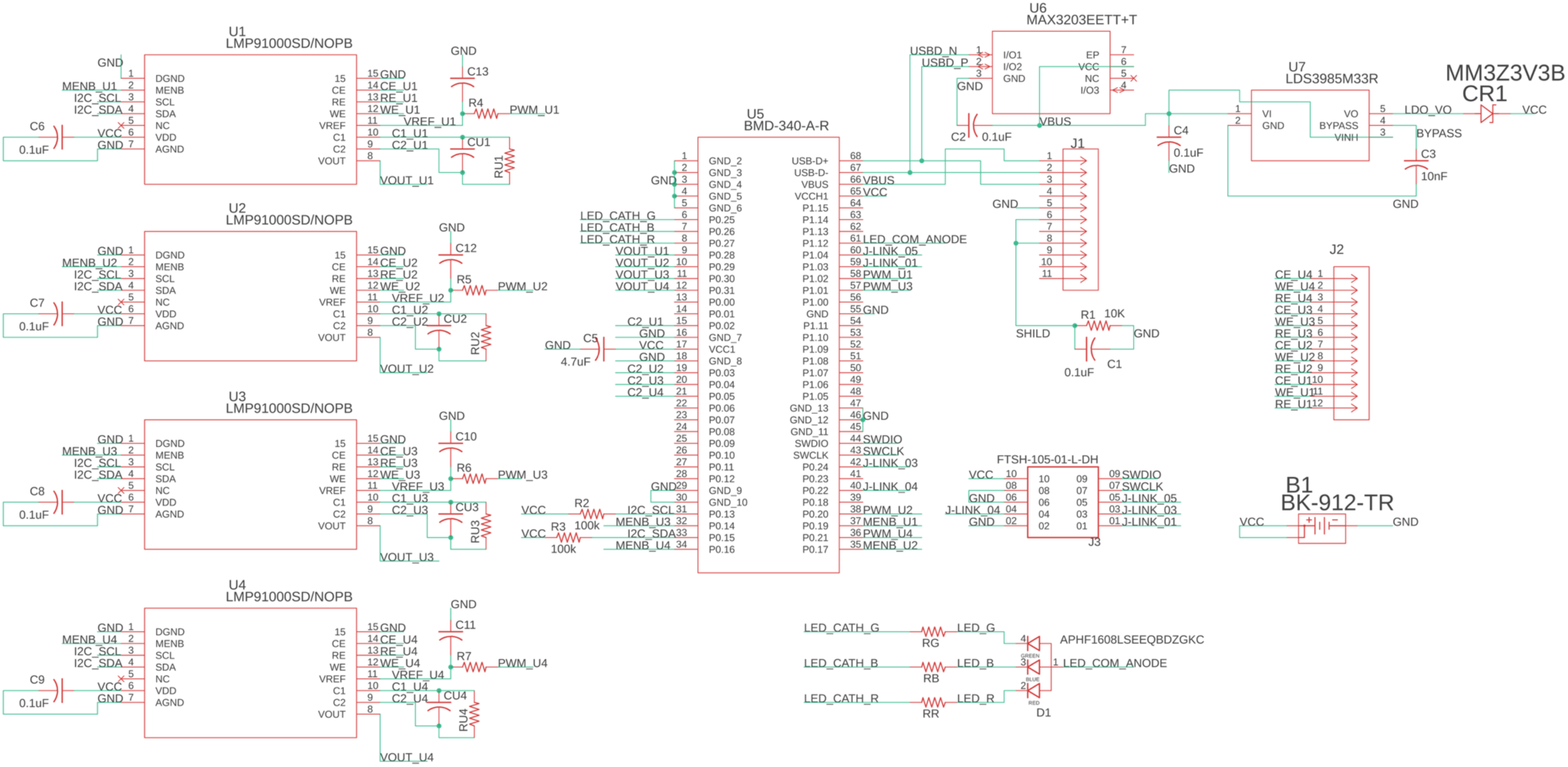
Circuit Schematic of the Device PCB.

